# Ligand Gaussian accelerated molecular dynamics (LiGaMD): Characterization of ligand binding thermodynamics and kinetics

**DOI:** 10.1101/2020.04.20.051979

**Authors:** Yinglong Miao, Apurba Bhattarai, Jinan Wang

## Abstract

Calculations of ligand binding free energies and kinetic rates are important for drug design. However, such tasks have proven challenging in computational chemistry and biophysics. To address this challenge, we have developed a new computational method “LiGaMD”, which selectively boosts the ligand non-bonded interaction potential energy based on the Gaussian accelerated molecular dynamics (GaMD) enhanced sampling technique. Another boost potential could be applied to the remaining potential energy of the entire system in a dual-boost algorithm (LiGaMD_Dual) to facilitate ligand binding. LiGaMD has been demonstrated on host-guest and protein-ligand binding model systems. Repetitive guest binding and unbinding in the β-cyclodextrin host were observed in hundreds-of-nanosecond LiGaMD simulations. The calculated binding free energies of guest molecules with sufficient sampling agreed excellently with experimental data (< 1.0 kcal/mol error). In comparison with previous microsecond-timescale conventional molecular dynamics simulations, accelerations of ligand kinetic rate constants in LiGaMD simulations were properly estimated using Kramers’ rate theory. Furthermore, LiGaMD allowed us to capture repetitive dissociation and binding of the benzamidine inhibitor in trypsin within 1 μs simulations. The calculated ligand binding free energy and kinetic rate constants compared well with the experimental data. In summary, LiGaMD provides a promising approach for characterizing ligand binding thermodynamics and kinetics simultaneously, which is expected to facilitate computer-aided drug design.

## Introduction

Free energy calculations of protein-ligand binding have been central to computational chemistry and biophysics research for the last several decades^1^. This is largely motivated by designing potent drug molecules with high binding free energies in the pharmaceutical industry^2^. A number of computational methods that have been developed in the field include thermodynamic integration (TI)^3^, free energy perturbation (FEP)^4^, double decoupling method^5^, umbrella sampling^6^, steered molecular dynamics (SMD)^7^, funnel metadynamics^8^, molecular mechanics Poisson-Boltzmann surface area and generalized Born surface area (MM/PBSA and MM/GBSA)^9^, and so on. Ligand binding free energy calculations have also been the focus of Statistical Assessment of Modeling of Proteins and Ligands (SAMPL)^10^ and Drug Design Data Resource (D3R)^11^ community challenges.

Kinetics of ligand binding have recently been recognized to be potentially more relevant for drug design. In particular, the dissociation rate constant that determines the drug residence time appears to better correlate with drug efficacy than the binding free energy^12^. However, ligand kinetic rates have proven even more difficult to compute than the binding free energies, largely due to slow processes of ligand binding and dissociation over long timescales^12b^. With remarkable advances in supercomputing and method developments, molecular dynamics (MD) simulations using specialized hardware^13^, Markov state model (MSM)^14^ and enhanced sampling^15^ have extended our capabilities to capture spontaneous ligand binding to target proteins. A number of enhanced sampling methods, including metadynamics^16^, random acceleration MD^17^, SMD^18^, weighted ensemble^19^ and selectively scaled MD^20^ have also been able to dissociate ligand molecules from the proteins. However, it remains extremely challenging to simulate repetitive ligand binding and dissociation processes, precluding accurate calculations of ligand kinetic rate constants.

With relatively small size and reduced complexity, host-guest binding systems has served as models of protein-ligand binding in SAMPL challenges^10^. The studied hosts include the cucurbit[7]uril (CB7), octa-acid, and CBClip and cyclodextrin (CD). These hosts are compounds that are typically much smaller than proteins but still large enough to bind guest molecules through non-covalent interactions, which share common characteristics as protein-ligand binding. Host-guest systems thus represent model systems for testing and improving free energy calculation methods. In addition to thermodynamics, Chang and co-workers have investigated kinetics of fast guest binding and dissociation in the β-CD host^21^. Tens to hundreds of repetitive guest binding and dissociation events were observed in microsecond-timescale MD simulations, which enabled comprehensive characterization of both thermodynamics and kinetics of the host-guest binding.

The MSM^22^ and metadynamics^8, 23^ have been applied to investigate the thermodynamics and kinetics of protein-ligand binding, using particularly the benzamidine inhibitor binding to trypsin as a model system. Multiple metadynamics trajectories with a total of 5 μs simulation time were obtained to predict the ligand unbinding pathways and dissociation rate constant, *k*_off_. The calculated *k*_off_ = 9.1 ± 2.5 s^-1^ was suggested to be in agreement of the experimental value 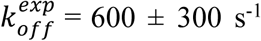. Separate funnel metadynamics simulations also allowed calculations of ligand binding free energies, in particular -8.5 ± 0.7 kcal/mol for the trypsin-benzamidine system^8^. The MSM built with 150 μs MD simulation data provided a complex picture of ligand binding kinetics and protein conformational plasticity^22^. The simulation predicted *k*_off_ = 131 ± 109 × 10^2^ s^-1^ appeared to be faster than the experimental value, while the predicted binding rate constant *k*_on_ = 6.4 ± 1.6 × 10^7^ M^-1^·s^-1^ compared well with the experimental value of 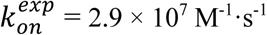. Repetitive ligand binding and unbinding in trypsin were also captured using a selective integrated-tempering-sampling molecular dynamics (SITSMD) method recently^24^. The binding free energy of benzamidine in trypsin was estimated as -5.06 kcal/mol, but the ligand kinetic rates were not calculated from the SITSMD simulations.

Here, we present a new computational method called ligand Gaussian accelerated molecular dynamics (“LiGaMD”), which enables us to simulate repetitive ligand binding and unbinding, and thus characterize both thermodynamics and kinetics of ligand binding simultaneously. Gaussian accelerated molecular dynamics (GaMD) is an enhanced sampling computational technique that works by adding a harmonic boost potential to smooth the biomolecular potential energy surface.^25^ GaMD greatly reduces energy barriers and accelerates biomolecular simulations by orders of magnitude.^26^ GaMD does not require pre-defined collective variables or reaction coordinates. Compared with the enhanced sampling methods that rely on careful selection of the collective variables, GaMD is of particular advantage for studying complex biological processes such as ligand binding to proteins^27^. Moreover, because the boost potential follows a Gaussian distribution, biomolecular free energy profiles can be properly recovered through cumulant expansion to the second order.^25^ GaMD builds on the previous accelerated MD (aMD) method^28^, but solves its energetic reweighting problem^29^ for free energy calculations of large biomolecules. GaMD has been implemented in the widely used AMBER^25, 30^, NAMD^31^ and GENESIS^32^ packages. GaMD has successfully revealed physical pathways and mechanisms of protein folding and ligand binding, which are consistent with experiments and long-timescale conventional MD (cMD) simulations.^25, 31, 33^ It has also been applied to characterize protein-protein^34^, protein-membrane,^35^ and protein-nucleic acid^36^ interactions.

Building upon GaMD, we have developed LiGaMD for more efficient sampling simulations of protein-ligand binding and unbinding processes. In LiGaMD, the non-bonded electrostatic and van der Waals interactions between the bound ligand and protein/environment are selectively boosted to enable ligand dissociation. In this context, selective acceleration has been found useful in previous enhanced sampling techniques, including the selective aMD^37^, selectively scaled MD^20^, essential energy space random walk^38^, replica exchange solute tempering (REST)^39^ and REST2^40^. In addition, a number of unbound ligand molecules are added in the solvent and another boost potential is applied on these ligand molecules, the protein and solvent in a dual-boost LiGaMD (LiGaMD_Dual) algorithm to facilitate ligand rebinding.

The LiGaMD method has been demonstrated on host-guest and protein-ligand binding model systems. Repetitive guest binding and unbinding in the β-CD host have been observed in hundreds-of-nanosecond LiGaMD simulations. The LiGaMD predicted guest binding free energies and kinetic rate constants agree well with those from previous cMD simulations^21^ and experimental data. Furthermore, LiGaMD has also allowed us to capture multiple dissociation and rebinding events of the benzamidine inhibitor in trypsin within 1 μs simulations. The reweighted ligand binding free energy and kinetic rate constants compared well with the experimental data.

## Methods

### Ligand Gaussian Accelerated Molecular Dynamics (LiGaMD)

Gaussian accelerated molecular dynamics (GaMD) is an enhanced sampling technique that works by adding a harmonic boost potential to smooth biomolecular potential energy surface and reduce the system energy barriers^25^. Details of the GaMD method have been described in previous studies^25, 31, 33a^. A brief summary is provided in **Appendix A**. Here, we develop a new ligand GaMD (LiGaMD) method for more efficient sampling of protein-ligand binding.

We consider a system of ligand *L* binding to a protein *P* in a biological environment *E*. The system comprises of *N* atoms with their coordinates 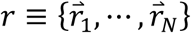 and momenta 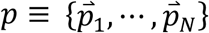. The system Hamiltonian can be expressed as:

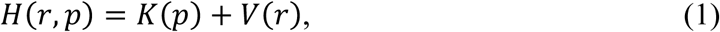

where *K*(*p*) and *V*(*r*) are the system kinetic and total potential energies, respectively. Next, we decompose the potential energy into the following terms:

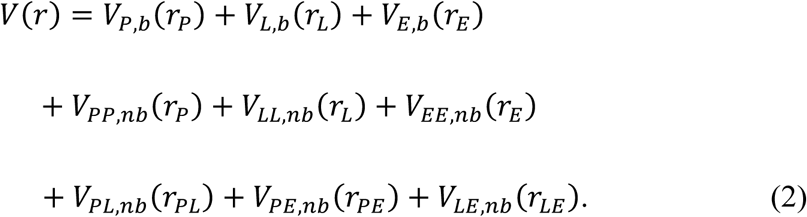

where *V*_*P,b*_, *V*_*L,b*_ and *V*_*E,b*_ are the bonded potential energies in protein *P*, ligand *L* and environment *E*, respectively. *V*_*PP,nb*_, *V*_*LL,nb*_ and *V*_*EE,nb*_ are the self non-bonded potential energies in protein *P*, ligand *L* and environment *E*, respectively. *V*_*PL,nb*_, *V*_*PE,nb*_ and *V*_*LE,nb*_ are the non-bonded interaction energies between *P-L, P-E* and *L-E*, respectively. According to classical molecular mechanics force fields^41^, the non-bonded potential energies are usually calculated as:

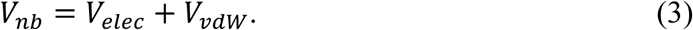

Where *V*_*elec*_ and *V*_*vdw*_ are the system electrostatic and van der Waals potential energies. Presumably, ligand binding mainly involves the non-bonded interaction energies of the ligand, *V*_*L,nb*_(*r*) = *V*_*LL,nb*_(*r*_*L*_) + *V*_*PL,nb*_(*r*_*PL*_) + *V*_*LE,nb*_(*r*_*LE*_). In LiGaMD, we add a boost potential selectively to the ligand non-bonded potential energy as:

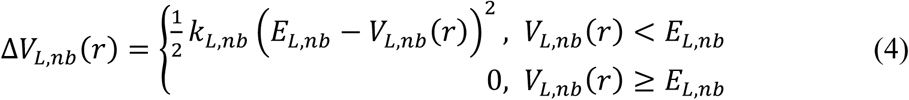

where *E*_*L,nb*_ is the threshold energy for applying boost potential and *k*_*L,nb*_ is the harmonic constant. For simplicity, the subscript of Δ*V*_*L,nb*_(*r*), *E*_*L,nb*_ and *k*_*L,nb*_ is dropped in the following. The LiGaMD simulation parameters are derived similarly as in the previous GaMD algorithm (**Appendix A**). When *E* is set to the lower bound *E=V*_*max*_, *k*_0_ can be calculated as:

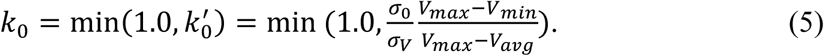

Alternatively, when the threshold energy *E* is set to its upper bound 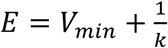, *k*_0_ is set to:

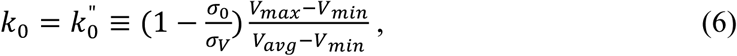

if 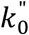 is found to be between *0* and *1*. Otherwise, *k*_0_ is calculated using Eqn. (5).

Next, one can add multiple ligand molecules in the solvent to facilitate ligand binding to proteins in MD simulations^13a, 42^. This is based on the fact that the ligand binding rate constant *k*_on_ is inversely proportional to the ligand concentration. The higher the ligand concentration, the faster the ligand binds, provided that the ligand concentration is still within its solubility limit. In addition to selectively boosting the bound ligand to accelerate its dissociation, another boost potential could thus be applied on the unbound ligand molecules, protein and solvent to facilitate ligand rebinding. The second boost potential is calculated using the total system potential energy other than the non-bonded potential energy of the bound ligand as:

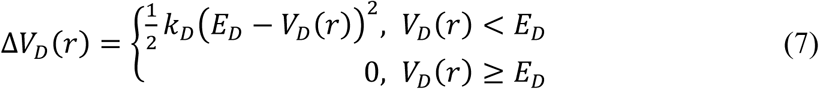

where *E*_*D*_ and *k*_*D*_ are the corresponding threshold energy for applying the second boost potential and the harmonic constant, respectively. This leads to dual-boost LiGaMD (LiGaMD_Dual) with the total boost potential Δ*V*(*r*) = Δ*V*_*L,nb*_(*r*) + Δ*V*_*D*_(*r*).

### Ligand Binding Free Energy Calculations from 3D Potential of Mean Force

We calculate ligand binding free energy from 3D potential of mean force (PMF) of ligand displacements from the target protein as the following^43^:

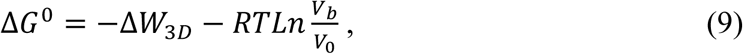

where *V*_0_ is the standard volume, *V*_*b*_ = ∫_+_ *e*^−*βW*(*r*)^ *dr* is the average sampled bound volume of the ligand with *β* = 1/*k*_*B*_*T, k*_*B*_ is the Boltzmann constant, *T* is the temperature, and Δ*W*_3*D*_ is the depth of the 3D PMF. Δ*W*_3*D*_ can be calculated by integrating Boltzmann distribution of the 3D PMF *W*(*r*) over all system coordinates except the x, y, z of the ligand:

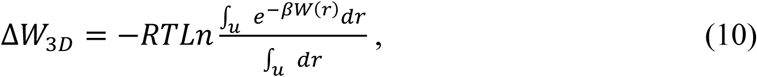

where *V*_*u*_ = ∫_*u*_ *dr* is the sampled unbound volume of the ligand. The exact definitions of the bound and unbound volumes *V*_*b*_ and *V*_*u*_ are not important as the exponential average cut off contributions far away from the PMF minima^43b^. A python script PyReweighting-3D.py is freely available in the PyReweighting tool kit (http://miao.compbio.ku.edu/PyReweighting/) for calculating the 3D PMF and associated ligand binding free energies. It works for both cMD (without energetic reweighting) and enhanced sampling simulations using LiGaMD with energetic reweighting (**Appendix B**).

### Ligand Binding Kinetics obtained from Reweighting of LiGaMD Simulations

Provided sufficient sampling of repetitive ligand dissociation and binding in simulations, one can record the time periods and calculate their averages for the ligand found in the bound (τ_*B*_) and unbound (τ_*U*_) states from the simulation trajectories. The τ_*B*_ corresponds to residence time in drug design^44^. Then the ligand dissociation and binding rate constants (*k*_off_ and *k*_on_) were calculated as:

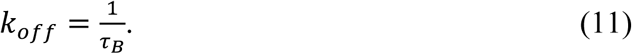

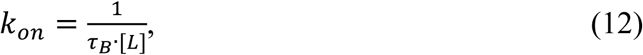

where [L] is the ligand concentration in the simulation system.

Two algorithms were implemented using the transition state theory (TST) and Kramers’ rate theory for reweighting kinetics of the LiGaMD simulations. According to Kramers’ Rate Theory (**Appendix C**), the rate of a chemical reaction in the large viscosity limit is calculated as^26^:

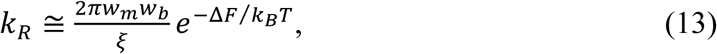

where *w*_*m*_ and *w*_*b*_ are frequencies of the approximated harmonic oscillators (also referred to as curvatures of free energy surface^45^) near the energy minimum and barrier, respectively, *ξ* is the apparent friction coefficient and Δ*F* is the free energy barrier of transition. The apparent friction coefficient *ξ* is related to the diffusion coefficient *D* with *ξ* = *k*_*B*_*T*/*D*. The apparent diffusion coefficient *D* can be obtained by dividing the kinetic rate calculated directly using the transition time series collected directly from simulations by that using the probability density solution of the Smoluchowski equation^46^ (**Appendix C**). In order to reweight ligand kinetics from the LiGaMD simulations using the Kramer’s rate theory, the free energy barriers of ligand binding and dissociation are calculated from the original (reweighted, **Δ*F***) and modified (no reweighting, **Δ*F\****) PMF profiles, similarly for curvatures of the reweighed (*w*) and modified (*w*^∗^, no reweighting) PMF profiles near the guest bound (“B”) and unbound (“U”) low-energy wells and the energy barrier (“Br”), and the ratio of apparent diffusion coefficients from LiGaMD simulations without reweighting (modified, *D*^∗^) and with reweighting (*D*). The resulting numbers are then plugged into Eq. (11) to estimate accelerations of the ligand binding and dissociation rates during LiGaMD simulations (**Appendix C**)^26^, which allows us to recover the original kinetic rate constants.

In comparison, the rate of a chemical reaction in the transition state theory (TST) is calculated as^47^:

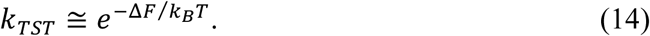

Only the energy barriers of ligand binding and dissociation need to be calculated from the original (reweighted, **Δ*F***) and modified (no reweighting, **Δ*F\****) free energy profiles for estimating the accelerations and recovering the ligand kinetic rate constants.

### Host-Guest Binding Simulations

GaMD simulations were performed on the binding of two guest molecules (aspirin and 1-butanol) to the β-CD host using the same input files as in a previous study by Tang and Chang^21^. The CD host were modeled with both the GAFF and q4MD force fields. The guest molecules were modeled with GAFF. The LiGaMD and dual-boost LiGaMD (LiGaMD_Dual) simulations were compared with GaMD simulations using the previous algorithms^25, 33b^, including the total potential boost GaMD (GaMD_Tot), dual-boost GaMD (GaMD_Dual)^25^, non-bonded potential boost GaMD (GaMD_NB) and non-bonded dual-boost GaMD (GaMD_Dual_NB)^33b^ simulations, as well as the previous long microsecond-timescale cMD simulations^21^ (**Tables 1 and S1**).

**Table 1.** Summary of host-guest binding thermodynamics and kinetics obtained from GaMD simulations on the binding of two guest molecules (aspirin and 1-butanol) to the β-cyclodextrin (CD) host. The CD host is modeled with the GAFF and q4MD force fields. *ΔV* is the system boost potential applied in GaMD simulations. *ΔG* is the ligand binding free energy. *N*_*D*_ and *N*_*B*_ are the number of host-guest dissociation and binding events collected from the individual simulations. *k*_*on*_*\** and *k*_*off*_*\** are the kinetic dissociation and binding rate constants, which are calculated from only the GaMD simulations with *N*_*D*_ > 0 and *N*_*B*_ > 0.

| Host | Ligand | GaMD<br>(300 ns x 3) | $\Delta V$<br>(kcal/mol) | $N_D$ | $N_B$ | $\Delta G$<br>(kcal/mol) | $k_{on}^*$<br>($\times 10^8 \text{ M}^{-1} \cdot \text{s}^{-1}$) | $k_{off}^*$<br>($\times 10^6 \text{ s}^{-1}$) |
| --- | --- | --- | --- | --- | --- | --- | --- | --- |
| CD:<br>GAFF | Aspirin | GaMD Tot | $6.48 \pm 2.59$ | 0, 2, 1 | 0, 1, 0 | - | - | - |
| | | GaMD Dual | $10.37 \pm 2.98$ | 0, 3, 0 | 0, 3, 0 | - | - | - |
| | | GaMD NB | $4.15 \pm 2.26$ | 1, 1, 2 | 1, 0, 1 | - | - | - |
| | | GaMD NB Dual | $8.75 \pm 2.82$ | 0, 1, 0 | 0, 1, 0 | - | - | - |
| | | LiGaMD | $2.60 \pm 1.27$ | 3, 1, 2 | 2, 1, 1 | $-4.90 \pm 0.03$ | $3.05 \pm 0.92$ | $40.67 \pm 24.25$ |
| | | LiGaMD Dual | $9.18 \pm 2.84$ | 2, 4, 2 | 2, 3, 2 | $-2.89 \pm 0.18$ | $3.93 \pm 0.98$ | $275.6 \pm 57.5$ |
| | 1-Butanol | LiGaMD | $2.54 \pm 1.00$ | 1, 0, 0 | 1, 0, 0 | - | - | - |
| | | LiGaMD Dual | $9.51 \pm 2.85$ | 1, 0, 1 | 1, 0, 1 | - | - | - |
| CD:<br>q4MD | Aspirin | GaMD Tot | $6.22 \pm 2.52$ | 0, 2, 0 | 0, 2, 0 | - | - | - |
| | | GaMD Dual | $10.77 \pm 3.02$ | 1, 0, 0 | 1, 0, 0 | - | - | - |
| | | GaMD NB | $4.13 \pm 2.17$ | 0, 2, 2 | 0, 2, 2 | - | - | - |
| | | GaMD NB Dual | $8.78 \pm 2.83$ | 0, 2, 2 | 0, 2, 2 | - | - | - |
| | | LiGaMD | $2.84 \pm 1.34$ | 4, 3, 2 | 3, 2, 2 | $-4.74 \pm 0.53$ | $8.99 \pm 5.02$ | $42.82 \pm 15.41$ |
| | | LiGaMD Dual | $10.06 \pm 3.15$ | 8, 7, 4 | 8, 6, 3 | $-3.34 \pm 0.15$ | $10.37 \pm 2.60$ | $56.61 \pm 19.05$ |
| | 1-Butanol | LiGaMD | $2.87 \pm 1.12$ | 9, 4, 7 | 9, 4, 7 | $-2.84 \pm 0.58$ | $8.76 \pm 2.93$ | $482.8 \pm 246.3$ |
| | | LiGaMD Dual | $9.74 \pm 2.93$ | 9, 13, 8 | 9, 13, 8 | $-1.34 \pm 0.10$ | $13.53 \pm 2.15$ | $436.8 \pm 90.7$ |

With the same restart files as used for running the cMD production simulations^21^, the GaMD simulations proceeded with 1 ns short cMD to collect potential statistics, 20 ns GaMD equilibration after adding the boost potential and then three independent 300 ns production runs. GaMD production frames were saved every 0.1 ps for analysis. The VMD^48^ and CPPTRAJ^49^ tools were used for simulation analysis. The number of host-guest dissociation and binding events (*N*_*D*_ and *N*_*B*_) and the guest binding and unbinding time periods (τ_*B*_ and τ_*U*_) were recorded from individual simulations (**Table S2**). With high fluctuations, τ_*B*_ and τ_*U*_ were recorded for only the time periods longer than 1 ns as applied in analysis of the previous cMD simulations^21^. For systems with more than one time of guest dissociation and binding in each of the individual simulations, 1D, 2D and 3D PMF profiles, as well as the host-guest binding free energies, were calculated through energetic reweighting of the GaMD simulations. The center-of-mass distances between the host and guests (d_HG_) and the host radius of gyration (*R*_*g*_) were chosen as reaction coordinates for calculating the 1D PMF profiles. 2D PMF profiles of (d_HG_, *R*_*g*_) were also calculated to analyze conformational changes of the CD host upon guest binding. The bin size was set to 0.5 Å for the d_HG_ and 0.05 Å for the *R*_*g*_. The cutoff of the number of simulation frames in one bin for reweighting was set to 500 in 1D and 2D PMF calculations. The 3D PMF profiles of guest displacements from the CD host in the X, Y and Z directions were further calculated from the LiGaMD simulations. The bin sizes were set to 1 Å in the X, Y and Z directions. The cutoff for the number of simulation frames in one bin for reweighting the 3D PMF was set to 10 for LiGaMD simulations and 50 for LiGaMD_Dual simulations. The host-guest binding free energies (*ΔG*) were calculated using the reweighted 3D PMF profiles and compared with those from previous cMD simulations and experimental data^21^ (**Table S1**). In addition, accelerations of the ligand dissociation and binding rate constants (*k*_*on*_ and *k*_*off*_) in LiGaMD simulations were analyzed using the TST and Kramers’ rate theory as described above (**Table S3**).

### Protein-Ligand Binding Simulations

LiGaMD simulations using the dual-boost scheme were performed on benzamidine binding to the trypsin protein. The X-ray crystal structure of benzamidine-bound trypsin (PDB ID: 3PTB^50^) was used with the calcium ion and water molecules kept. The missing hydrogen atoms were added using by the *tleap* module in AMBER^51^. The general Amber force field^52^ and the AMBER ff14SB force field^53^ were used for the ligand and protein, respectively. Atomic partial charges of benzamidine were obtained through B3LYP/6-31G* quantum calculations of the electrostatic potential, for which the RESP charges^54^ were fitted using the antechamber program^51^. The system was neutralized by adding a number of counter ions (Cl^-^) and immersed in a cubic TIP3P water ^55^box, which was extended 13 Å from the protein-ligand complex. Testing simulations were performed on protonation of the protein active-site residue His57 at the N_δ_ atom or the N_ε_ atom. Results showed that benzamidine could bind to the protein target site as in the X-ray conformation with the N_δ_ atom protonated, but not with atom N_ε_ protonated. Therefore, the protein residue His57 was protonated at the N_δ_ atom. Moreover, a total of 10 ligand molecules (one in the X-ray bound conformation and another nine placed randomly in the solvent) were included in the system to facilitate ligand binding. This design was based on the fact that the ligand binding rate constant *k*_on_ is inversely proportional to the ligand concentration. The higher the ligand concentration, the faster the ligand binds, provided that the ligand concentration is still within its solubility limit.

The built simulation system was energy minimized with 1 kcal/mol/Å^2^ constraints on the heavy atoms of the protein and ligand, including the steepest descent minimization of 5,000 steps followed by a conjugate gradient minimization of 5,000 steps. The system was then heated from 0 K to 300 K for 200 ps. It was further equilibrated using the NVT ensemble at 300K for 800 ps and the NPT ensemble at 300 K and 1 bar for 1 ns with 1 kcal/mol/Å^2^ constraints on the heavy atoms of the protein and ligand, followed by 2 ns short cMD without any constraint. The LiGaMD simulations proceeded with 14 ns short cMD to collect the potential statistics, 54.6 ns GaMD equilibration after adding the boost potential and then five independent 1000 ns production runs. Initial testing simulations showed that when the threshold energy for applying boost potential to the ligand non-bonded energy was set to the lower bound (i.e., *E* = *V*_max_), the bound ligand maintained the X-ray conformation and did not dissociate. In comparison, when the threshold energy was set to the upper bound (i.e., *E* = *V*_min_+1/*k*), it enabled high enough boost potential to dissociate the ligand from the protein. Therefore, the threshold energy for applying the ligand boost potential was set to the upper bound in the LiGaMD_Dual simulations. For the second boost potential that was applied to the system total potential energy other than the ligand non-bonded potential energy, sufficient acceleration was obtained to sample ligand rebinding by setting the threshold energy to the lower bound. LiGaMD_Dual production simulation frames were saved every 0.2 ps for analysis.

The VMD^48^ and CPPTRAJ^49^ tools were used for simulation analysis. The number of ligand dissociation and binding events (*N*_*D*_ and *N*_*B*_) and the ligand binding and unbinding time periods (τ_*B*_ and τ_*U*_) were recorded from individual simulations (**Table S4**). With high fluctuations, τ_*B*_ and τ_*U*_ were recorded for only the time periods longer than 5 ns. The 1D, 2D and 3D PMF profiles, as well as the ligand binding free energy, were calculated through energetic reweighting of the LiGaMD_Dual simulations. The distance between the N atom in benzamidine and CG atom of Asp189 in trypsin was chosen as the reaction coordinate for calculating the 1D PMF profiles. The bin size was set to 1.0 Å. 2D PMF profiles of the benzamidine:N – Asp189:CG and Trp215:NE – Asp189:CG atom distances were also calculated to analyze conformational changes of the protein upon ligand binding. The bin size was set to 1.0 Å for the atom distances. The cutoff for the number of simulation frames in one bin was set to 500 for reweighting 1D and 2D PMF profiles. The 3D PMF profiles of guest displacements from the CD host in the X, Y and Z directions were further calculated from the LiGaMD simulations. The bin sizes were set to 1 Å in the X, Y and Z directions. The cutoff of simulation frames in one bin for 3D PMF reweighting (ranging from 1100-4000 for five individual LiGaMD_Dual simulations) was set to the minimum number below which the calculated 3D PMF minimum will be shifted. The ligand binding free energies (*ΔG*) were calculated using the reweighted 3D PMF profiles. Furthermore, structural clustering was performed on frames of the diffusing ligand molecules from each 1 μs LiGaMD_Dual simulation trajectory using the Density Based Spatial Clustering of Applications with Noise (DBSCAN) algorithm^56^ in CPPTRAJ^49^. The frames were sieved at a stride of 500 for clustering. The remaining frames were assigned to the closest cluster afterwards. The distance cutoff for DBSCAN clustering was set to 0.5 Å. The resulting structural clusters were reweighted to obtain energetically significant binding pathways of the ligand. In addition, the ligand dissociation and binding rate constants (*k*_*on*_ and *k*_*off*_) were calculated from the LiGaMD_Dual simulations with their accelerations analyzed using the Kramers’ rate theory as described above (**Table S5**).

## Results

### Thermodynamics of Host-Guest Binding

For the β-cyclodextrin (CD) host as modeled with both the GAFF and q4MD force fields, all-atom GaMD simulations were performed to investigate the dissociation and binding of two guest molecules aspirin (**Fig. 1A**) and 1-butanol (**Fig. 1B**) using a number of different potential boost algorithms (**Table 1**). The center-of-mass distances between the host and guest molecules were monitored as a function of time to record the number of dissociation and binding events (*N*_*D*_ and *N*_*B*_) in each of the 300 ns GaMD simulations. The new LiGaMD method especially using the dual-boost algorithm (LiGaMD_Dual) showed significantly improved sampling of ligand binding compared with the previous GaMD algorithms, including the total potential boost GaMD (GaMD_Tot), dual-boost GaMD (GaMD_Dual), non-bonded potential boost GaMD (GaMD_NB) and non-bonded dual-boost GaMD (GaMD_Dual_NB) (**Table 1**). Repetitive binding and unbinding of guest molecules were observed in 300 ns LiGaMD and LiGaMD_Dual simulations, except for the binding of 1-butanol to CD that was modeled with the GAFF force field (**Fig. S1**). In comparison, no guest dissociation and/or binding was observed in one or more of the 300 ns GaMD simulations using the other algorithms (**Table 1**).

**Fig. 1.**
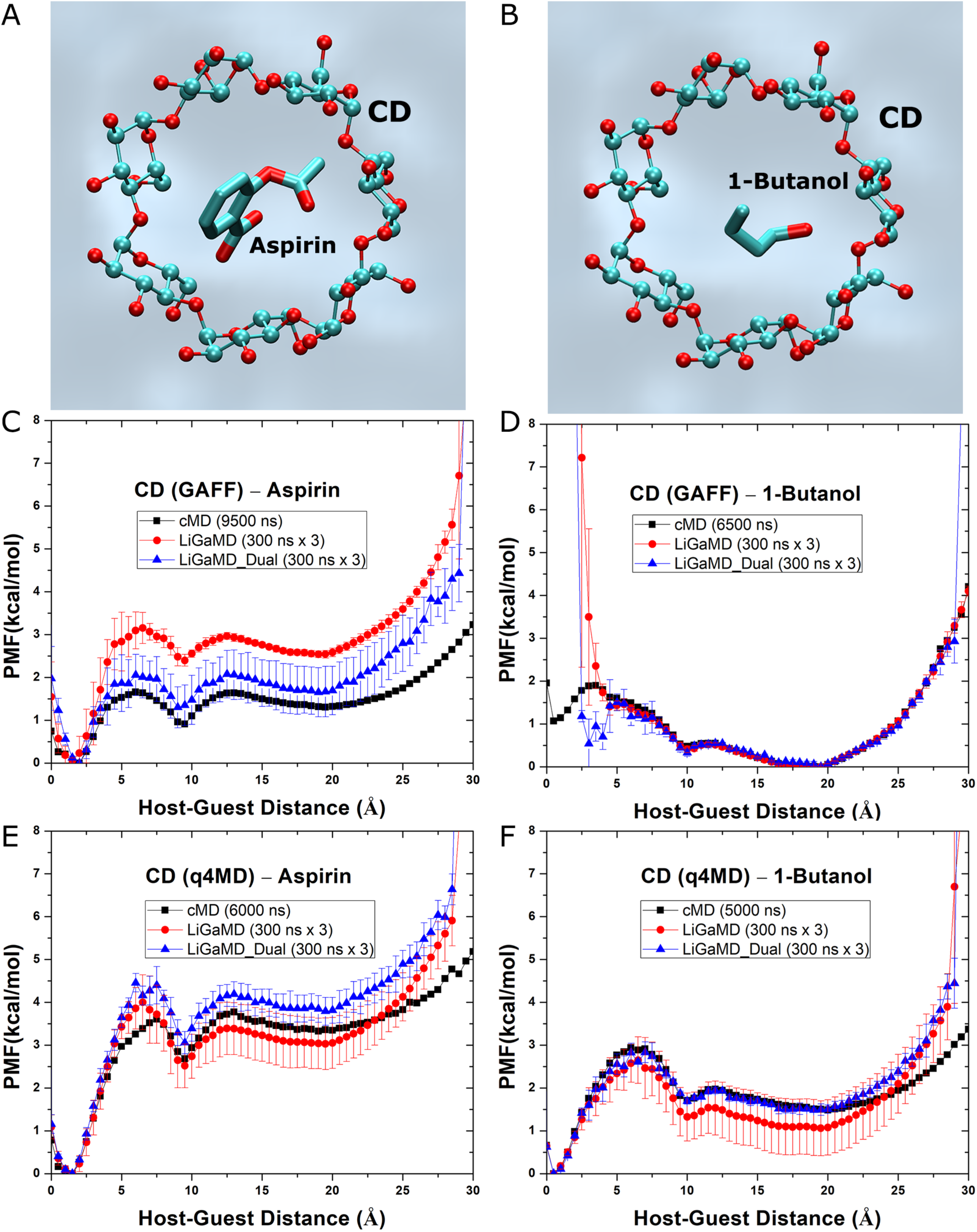
Comparison of potential of mean force (PMF) free energy profiles calculated from conventional MD (cMD), LiGaMD and dual-boost LiGaMD (LiGaMD_Dual) simulations of host-guest binding: (A-B) Computational models of the β-cyclodextrin (CD) host (balls-and-sticks) in the presence of guest (A) aspirin and (B) 1-butanol (thick sticks) in aqueous medium (cyan). (C-F) PMF profiles calculated from microsecond-timescale cMD simulations and three independent 300 ns LiGaMD and LiGaMD_Dual simulations of (C) CD using the GAFF force field with aspirin, (D) CD using the GAFF force field with 1-butanol, (E) CD using the q4MD force field with aspirin, (F) CD using the q4MD force field with 1-butanol.

Provided improved sampling in the LiGaMD and LiGaMD_Dual simulations, we computed potential of mean force (PMF) free energy profiles to characterize the host-guest binding quantitatively (**Fig. 1**). The host-guest distance (d_HG_) was chosen as a reaction coordinate. The PMF profiles calculated from three 300 ns LiGaMD or LiGaMD_Dual simulations were compared with those from microsecond-timescale cMD simulations of CD using the GAFF force field with aspirin (**Fig. 1C**), CD using the GAFF force field with 1-butanol (**Fig. 1D**), CD using the q4MD force field with aspirin (**Fig. 1E**), and CD using the q4MD force field with 1-butanol (**Fig. 1F**). For binding of 1-butanol to CD modeled with GAFF, while 300 ns LiGaMD and LiGaMD_Dual simulations sampled the system unbound (d_HG_ > ∼15 Å) and intermediate (d_HG_ = ∼10 Å) states, being closely similar to the 6500 ns cMD simulations, the bound state (d_HG_ = ∼0.5 Å) was poorly sampled in the LiGaMD simulations (**Fig. 1D**). This correlated with the above finding that the guest dissociation and binding were seldomly observed in these simulations (**Table 1** and **Figs. S1C-S1D**). For the other three systems, 300 ns LiGaMD and LiGaMD_Dual simulations sampled the same bound (d_HG_ = ∼0.5-2 Å), intermediate (d_HG_ = ∼10 Å) and unbound (d_HG_ > ∼15 Å) low-energy states as in the microsecond timescale cMD simulations (**Figs. 1C, 1E** and **1F**). Their global energy minima were all identified in the bound state. However, differences were found in the magnitudes of PMF profiles near the intermediate, unbound and energy barrier regions. Overall, the LiGaMD_Dual simulations showed better agreements with the long-timescale cMD simulations than LiGaMD, especially for binding of 1-butanol to CD modeled with q4MD (**Fig. 1F**). The LiGaMD_Dual provided the most efficient sampling of host-guest binding and dissociation events (**Table 1** and **Fig. S1**) and the closest free energy profiles as compared with the reference cMD simulations (**Fig. 1**).

Next, we analyzed conformational changes of the CD host modeled with GAFF (**Fig. 2**) and q4MD (**Fig. 3**) upon guest binding. PMF profiles were calculated for the radius of gyration (*R*_*g*_) of the CD host modeled with GAFF from the cMD simulations in the ligand-free (*apo*), aspirin and 1-butanol binding forms (**Fig. 2A**). Two low-energy states, including “Compact” (*R*_*g*_ = ∼5.7 Å) and “Open” (*R*_*g*_ = ∼5.9 Å), were identified for the CD host. While the *apo* and 1-butanol bound CD predominantly adopted the compact conformational state, binding of aspirin biased conformational ensemble of CD towards the open state. In this regard, aspirin (**Fig. 1A**) appeared to be larger in size than 1-butanol (**Fig. 1B**). Furthermore, we calculated 2D profiles of (d_HG_, *R*_*g*_) and identified low-energy conformational states of the system. For aspirin binding, three low-energy states were found from the 9500 ns cMD simulation, including the “Bound (B)”, “Intermediate (I)” and “Unbound (U)” states, in which the CD host adopted primarily the Open, Compact and Compact conformations, respectively (**Figs. 2B** and **2C**). The 2D PMF profile of aspirin binding calculated from three 300 ns LiGaMD_Dual simulations combined was highly similar to that from 9500 ns cMD simulation, depicting the same three low-energy conformational states (**Figs. 2C** and **2D**). In comparison, only the Bound and Unbound low-energy states were clearly identified for binding of 1-butanol in the 2D PMF profiles of both 6500 ns cMD (**Fig. 2E**) and three 300 ns LiGaMD_Dual simulations (**Fig. 2F**), while the intermediate state was hardly observed with apparently a very shallow energy well as shown in the 1D PMF profiles (**Fig. 1D**).

**Fig. 2.**
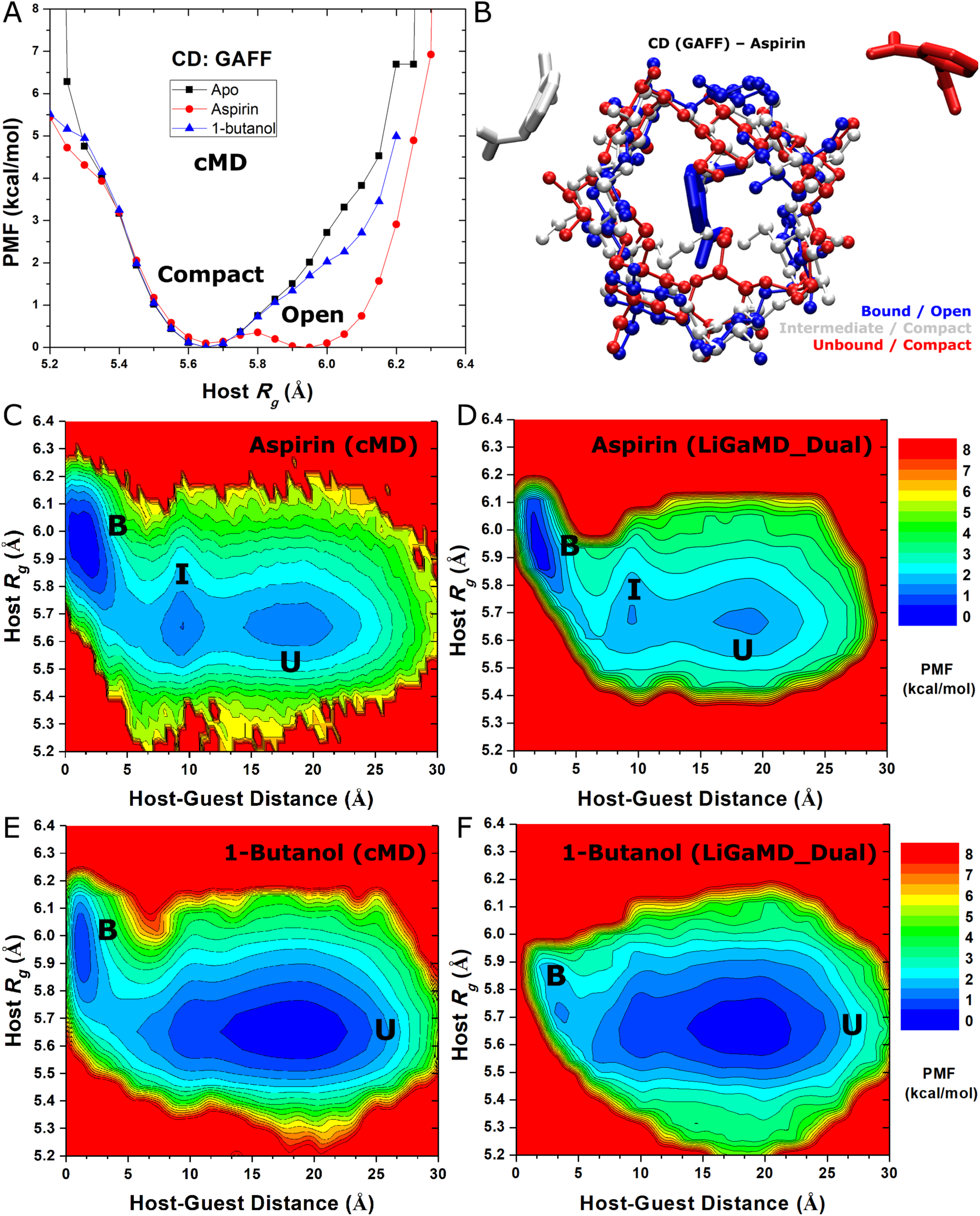
Free energy profiles and low-energy conformational states of guest binding to the CD host that was modeled with the GAFF force field: (A) PMF profiles of the host radius of gyration (*R*_*g*_) calculated from cMD simulations in the ligand-free (apo), aspirin and 1-butanol binding forms. Two low-energy states (“Compact” and “Open”) are identified for the CD host. (B) Three representative conformational states observed in simulations of guest aspirin binding to CD modeled with GAFF: the “Bound (B)”, “Intermediate (I)” and “Unbound (U)”, in which the CD host adopted primarily the Open, Compact and Compact conformations, respectively. (C-E) 2D PMF profiles of the host *R*_*g*_ versus the center-of-mass distance between host-guest of (C) aspirin binding from 9500 ns cMD simulation, (D) aspirin binding from three 300 ns LiGaMD_Dual simulation combined, (E) 1-butanol binding from 6500 ns cMD simulation, and (F) 1-butanol binding from three 300 ns LiGaMD_Dual simulation combined. The low-energy states are labeled.

**Fig. 3.**
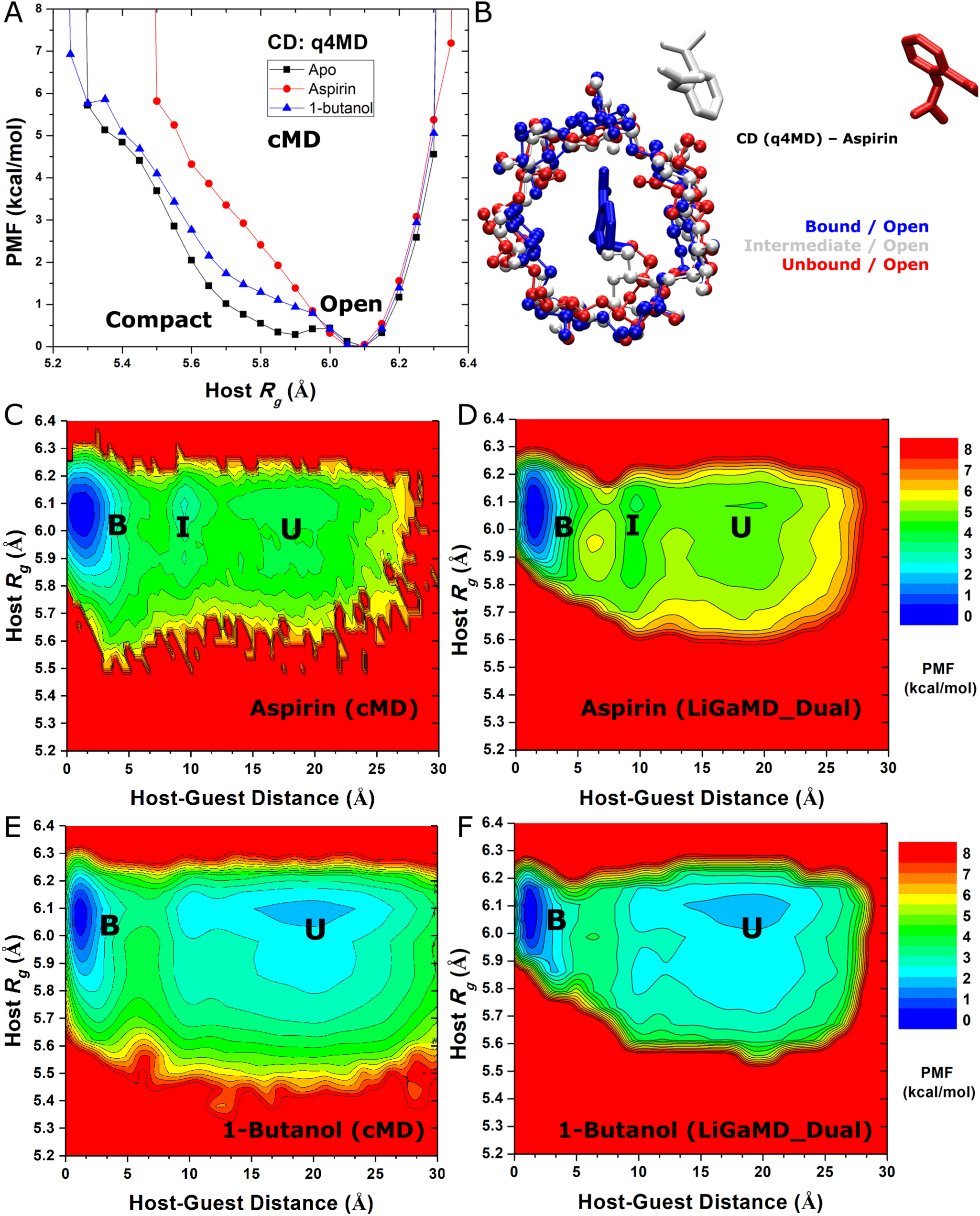
Free energy profiles and low-energy conformational states of guest binding to the CD host that was modeled with the q4MD force field: (A) PMF profiles of the host radius of gyration (*R*_*g*_) calculated from cMD simulations in the ligand-free (apo), aspirin and 1-butanol binding forms. Two low-energy states (“Compact” and “Open”) are labeled for the CD host. (B) Three representative conformational states observed in simulations of aspirin binding to CD modeled with q4MD: the “Bound (B)”, “Intermediate (I)” and “Unbound (U)”, in which the CD host all adopted primarily the Open conformation. (C-E) 2D PMF profiles regarding the host *R*_*g*_ versus center-of-mass distance between host-guest of (C) aspirin binding from 6000 ns cMD simulation, (D) aspirin binding from three 300 ns LiGaMD_Dual simulation combined, (E) 1-butanol binding from 5000 ns cMD simulation, and (F) 1-butanol binding from three 300 ns LiGaMD_Dual simulation combined. The low-energy states are labeled.

Furthermore, we evaluated the effects of different force field parameters for the CD host. With the q4MD force field, the CD host adopted predominantly the Open conformation with *R*_g_ = ∼6.1 Å in PMF profiles of both the *apo* and guest-bound forms, although it still sampled the Compact conformation with a shallow and broad energy well in the *apo* form (**Fig. 3A**). This was in contrast to the above findings that the CD host using the GAFF force field preferred the Compact conformation in the *apo* form, while guest binding especially aspirin induced the host to open (**Fig. 2**). Therefore, the usage of different force fields generated distinct structural dynamics and free energy profiles of host-guest binding. Nevertheless, consistent results were obtained from LiGaMD_Dual and cMD simulations provided the same system setup and force field. In this context, LiGaMD only enhanced conformational sampling of the studied systems, yielding similar results as obtained from significantly longer cMD simulations.

In 6000 ns cMD simulation of aspirin binding to CD modeled with q4MD, three low-energy conformational states were identified from 2D PMF profiles of the host-guest distance d_HG_ and host *R*_*g*_, including the “Bound (B)”, “Intermediate (I)” and “Unbound (U)” states, in which the CD host all adopted primarily the Open conformation (**Figs. 3B** and **3C**). A closely similar 2D PMF profile was obtained from three 300 ns LiGaMD_Dual simulations combined (**Figs. 3D**). Similar 2D profiles were obtained for 1-butanol binding from three 300 ns LiGaMD_Dual simulations combined and 5000 ns cMD simulation, during which CD was open in both the guest bound and unbound states (**Figs. 3E** and **3F**).

In addition to the 1D and 2D PMF profiles, we computed 3D PMF profiles of guest binding to the CD host in the X, Y and Z directions and then the guest binding free energies (see details in **Methods**). The host-guest binding free energies calculated from LiGaMD and LiGaMD_Dual simulations were compared with those obtained from previous cMD simulations and experimental data **(Tables 2** and **S1)**. Compared with experimental data, LiGaMD_Dual provided generally more accurate estimates of the guest binding free energies than LiGaMD (**Table 2**), being consistent with the finding that improved sampling with more dissociation and binding events was observed in the LiGaMD_Dual simulations (**Table 1**). For aspirin, the binding free energy errors were reduced from -1.16 ± 0.03 kcal/mol to 0.85 ± 0.18 kcal/mol with the CD host modeled from GAFF and from -1.00 ± 0.53 kcal/mol to 0.40 ± 0.15 kcal/mol with CD modeled using q4MD. For 1-butanol, the binding free energy error was calculated from only the simulations with CD modeled using q4MD, during each of which multiple dissociation and binding events were observed (**Table 1**). The binding free energy error of 1-butanol decreased from -1.17 ± 0.61 kcal/mol in LiGaMD simulations to 0.33 ± 0.21 kcal/mol in LiGaMD_Dual simulations (**Table 2**). The guest binding free energy errors from the LiGaMD_Dual simulations were less than 1 kcal/mol and mostly comparable to those from previous microsecond-timescale (5-9.5 μs) cMD simulations, despite variations in the guest binding free energies calculated using two different algorithms from the cMD simulations (*ΔG*_*comp1*_ and *ΔG*_*comp2*_) as adapted from Ref. ^**21**^.

**Table 2.** Comparison of the host-guest binding free energies calculated from the LiGaMD simulations, LiGaMD_Dual simulations, cMD simulations^**21**^ and experimental data^**21**^.

| Host | Ligand | $\Delta G_{sim} - \Delta G_{exp}$ (kcal/mol) | | | |
| --- | --- | --- | --- | --- | --- |
| | | cMD<br>($\Delta G_{comp1}$) | cMD<br>($\Delta G_{comp2}$) | LiGaMD | LiGaMD_Dual |
| CD:<br>GAFF | Aspirin | $-0.10 \pm 0.35$ | $1.47 \pm 0.06$ | $-1.16 \pm 0.03$ | $0.85 \pm 0.18$ |
| CD:<br>q4MD | Aspirin | $-2.59 \pm 0.37$ | $-0.37 \pm 0.05$ | $-1.00 \pm 0.53$ | $0.40 \pm 0.15$ |
| | 1-Butanol | $0.34 \pm 0.39$ | $-0.60 \pm 0.19$ | $-1.17 \pm 0.61$ | $0.33 \pm 0.21$ |

In summary, the PMF profiles and binding free energies of guest molecules in the CD host calculated from hundreds-of-nanosecond LiGaMD_Dual simulations agreed excellently with those from experimental data and previous microsecond-timescale cMD simulations^**21**^. Errors in the guest binding free energies were smaller than 1 kcal/mol from the LiGaMD_Dual simulations as compared with the experimental data. Therefore, both efficient enhanced sampling and accurate free energy calculations of host-guest binding were achieved through the LiGaMD_Dual simulations.

### Kinetics of Host-Guest Binding

In addition to the thermodynamic free energies, kinetic dissociation and binding rate constants were further derived from the relevant LiGaMD and LiGaMD_Dual simulations and compared with those from cMD simulations (**Table 3**). While the binding rate constants of guest molecules decreased to ∼30%-90% of those calculated from cMD simulations^**21**^, the dissociation rate constants increased by ∼1.7-15 times in LiGaMD simulations and ∼11-18 times in LiGaMD_Dual simulations (**Table 3**). In this context, long-timescale cMD simulations with repetitive host-guest binding and unbinding were available for comparison. However, in most protein-ligand binding studies (e.g., the trypsin-benzamidine binding described below) such cMD simulations are often not available, due to the extreme challenge of sampling free ligand binding and dissociation over long timescales. In this regard, we sought to reweight kinetics of LiGaMD and recover the original kinetic dissociation and binding rate constants from only the LiGaMD enhanced sampling simulations as follows.

**Table 3.**
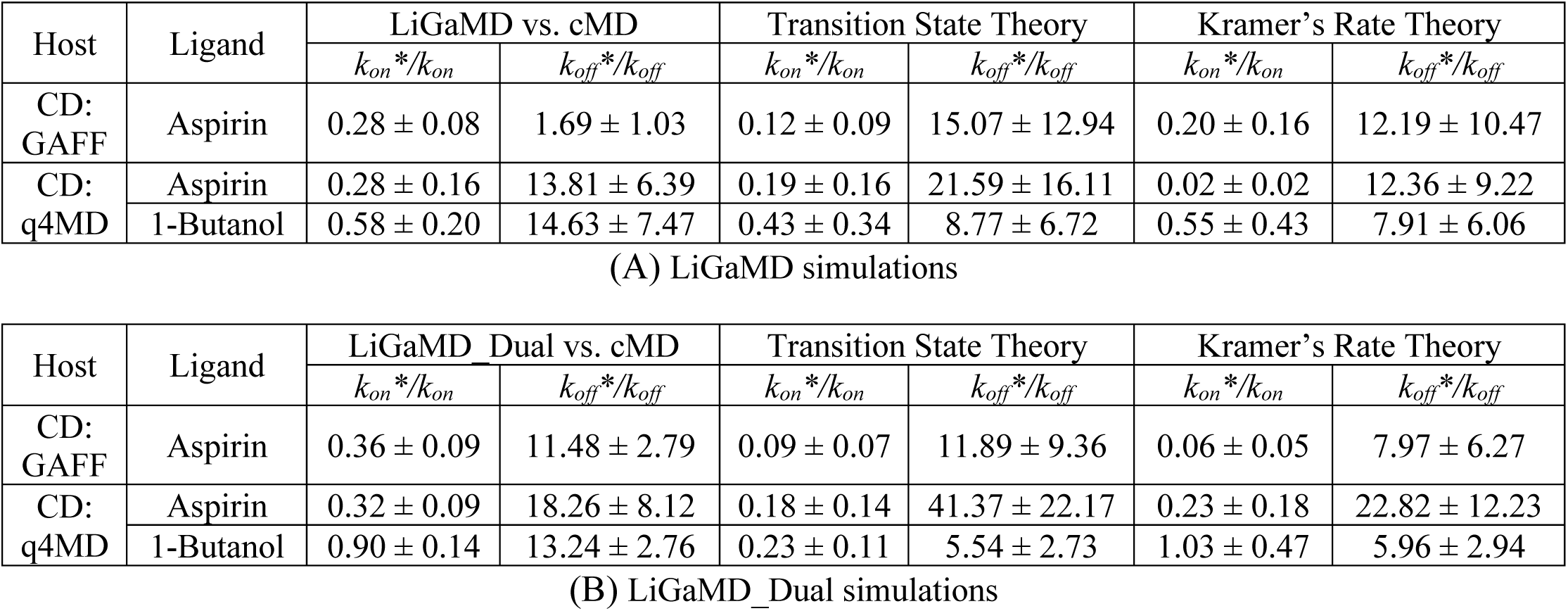
Accelerations of host-guest dissociation and binding rate constants obtained from (A) LiGaMD and (B) LiGaMD_Dual simulations as compared directly with previous cMD simulations^**21**^ and those derived using the Transition State Theory and Kramers’ Rate Theory.

Two algorithms were implemented using the transition state theory (TST) and Kramers’ rate theory for reweighting kinetics of the LiGaMD simulations (see details in **Methods**). In both algorithms, the energy barriers of ligand binding and dissociation were calculated from the original (reweighted) and modified (no reweighting) free energy profiles of the LiGaMD simulations. Since the reweighted PMF profiles were already obtained for the host-guest distances from the LiGaMD and LiGaMD_Dual simulations (**Fig. 1**), we also calculated the corresponding modified PMF profiles without energetic reweighting for comparative analysis (**Fig. 4**). Note that the bound state of 1-butanol in CD modeled with GAFF was poorly sampled in the LiGaMD and LiGaMD_Dual simulations (**Figs. 1D, S1** and **S2**). This system was thus excluded for both binding free energy and kinetics calculations. For the other three systems, the energy barriers were significantly reduced in the modified PMF profiles of both the LiGaMD and LiGaMD_Dual simulations for guest dissociation (**Fig. 4** and **Table S3**). In LiGaMD_Dual simulations of aspirin binding to CD modeled with GAFF, the dissociation free energy barrier (Δ*F*_*off*_) decreased by 73% from 2.06 ± 0.28 kcal/mol in the reweighted PMF profile to 0.59 ± 0.37 kcal/mol in the modified PMF profile (**Fig. 4B**). In the other LiGaMD and LiGaMD_Dual simulations, Δ*F*_*off*_ mostly decreased by ∼50% (**Fig. 4** and **Table S3**). Correspondingly, *k*_off_ estimated using the TST increased by ∼9-22 times in the LiGaMD simulations of aspirin and 1-butanol binding to CD and ∼6-41 times in the LiGaMD_Dual simulations (**Table 3**). On the other hand, the free energy barrier for ligand binding (Δ*F*_*on*_) actually increased in all modified PMF profiles of the LiGaMD and LiGaMD_Dual simulations than in the reweighted profiles (**Fig. 4** and **Table S3**). According the TST, the corresponding binding rate constant (*k*_*on*_) decreased by a factor ∼0.1-0.4 in LiGaMD simulations and ∼0.1-0.2 in LiGaMD_Dual simulations (**Table 3**). Despite the slower binding, significant accelerations were achieved in the rate-limiting dissociation of guest molecules during LiGaMD and LiGaMD_Dual simulations. Therefore, the guest dissociation and binding appeared to be more balanced in LiGaMD than in cMD, thereby resulting in overall improved sampling.

**Fig. 4.**
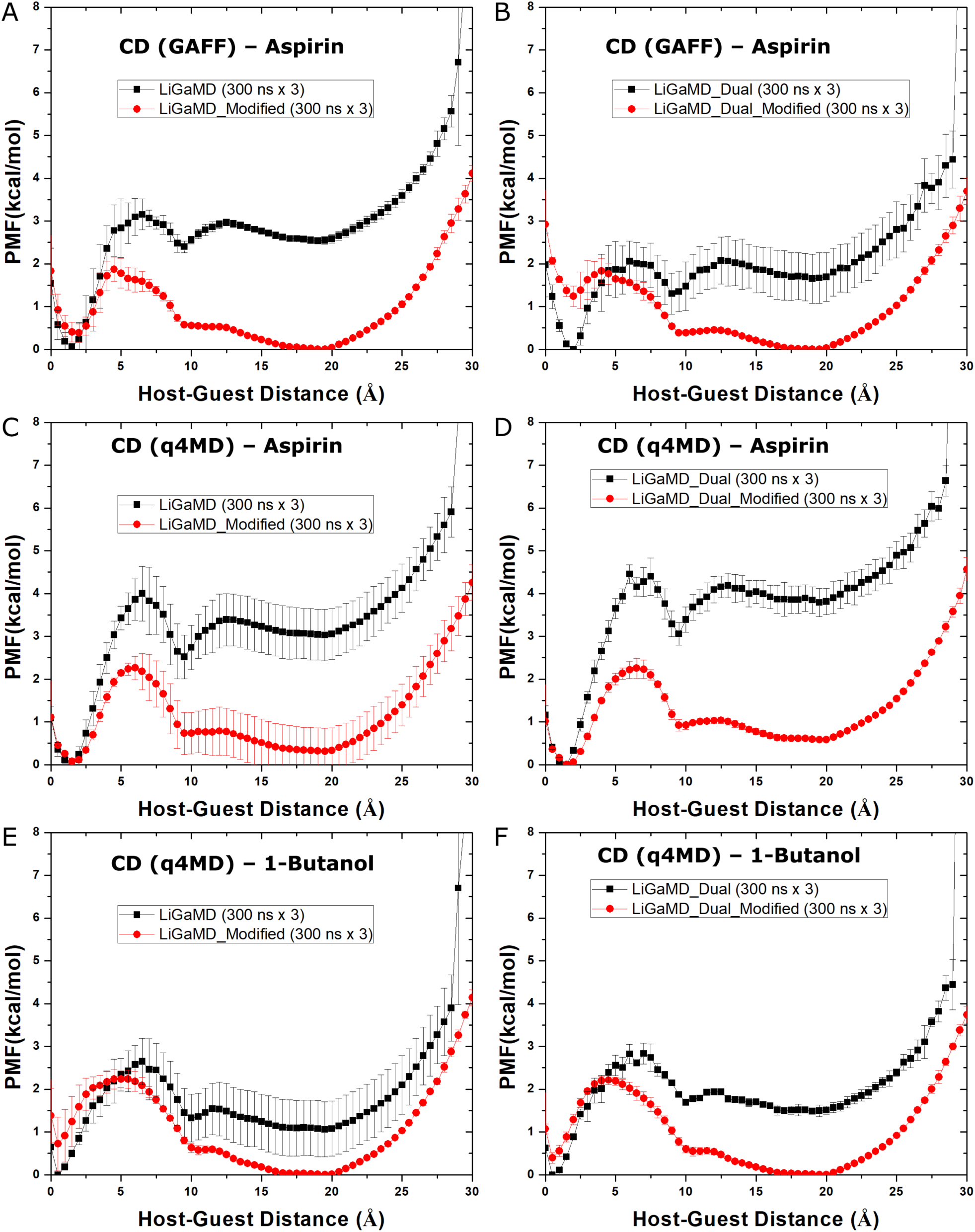
(A-F) The original (reweighted) and modified (no reweighting) PMF profiles of (A) LiGaMD simulations of CD using the GAFF force field with aspirin, (B) LiGaMD_Dual simulations of CD using the GAFF force field with aspirin, (C) LiGaMD simulations of CD using the q4MD force field with aspirin, (D) LiGaMD_Dual simulations of CD using the q4MD force field with aspirin, (E) LiGaMD simulations of CD using the q4MD force field with 1-butanol, and (F) LiGaMD_Dual simulations of CD using the q4MD force field with 1-butanol.

For applying the Kramers’ rate theory, curvatures of the reweighed (*w*) and modified (*w*^∗^, no reweighting) free energy profiles were further calculated near the guest bound (“B”) and unbound (“U”) energy minima and the energy barrier (“Br”), as well as the ratio of apparent diffusion coefficients calculated from the LiGaMD and LiGaMD_Dual simulations with reweighting (*D*) and without reweighting (modified, *D*^∗^) (**Table S3**). The resulting numbers were plugged into the Kramers’ rate equation to estimate accelerations of the guest binding and dissociation rate constants. Particularly, the *k*_off_ increased by ∼8-12 times in LiGaMD simulations and ∼6-23 times in LiGaMD_Dual simulations (**Table 3**). The *k*_on_ decreased by a factor of 0.02-0.55 in LiGaMD simulation. In LiGaMD_Dual simulations, *k*_on_ remained the same for 1-butanol binding to CD modeled with q4MD, but also decreased by factors of 0.06 and 0.23 for aspirin binding to CD modeled with GAFF and q4MD force fields, respectively (**Table 3**).

In summary, accelerations of host-guest dissociation and binding obtained from LiGaMD and LiGaMD_Dual simulations were derived using the TST and Kramers’ Rate Theory. Compared with cMD simulations, while the binding rates decreased to a certain extent, guest unbinding was significantly accelerated by ∼1.7-15 times in LiGaMD and 11-18 times in LiGaMD_Dual. Overall, the Kramers’ rate theory provided more accurate estimates of the ligand kinetic rate accelerations in the LiGaMD simulations than the TST (**Table 3**). It was thus applied for reweighting of ligand kinetics in analysis of further LiGaMD simulations.

### Thermodynamics of Ligand Binding to the Trypsin Model Protein

In addition to host-guest binding, LiGaMD was further tested on protein-ligand binding using trypsin as a model system. Initial testing simulations suggested the following setup for proper simulations of benzamidine binding to trypsin: First, residue His57 at the protein active site was protonated at the N_δ_ atom rather than the default N_ε_ atom, which was also depicted for the catalytic triad at the active site of proteases^57^. Testing simulations with the N_ε_ atom protonated in residue His57 showed that the benzamidine inhibitor could not bind to the protein target site as in the X-ray crystal conformation with ∼4.3 Å minimum root-mean square deviation (RMSD). In comparison, simulations with N_δ_ atom could capture repetitive binding of benzamidine to the target site with ∼1.0 Å minimum RMSD compared with the X-ray conformation (**Fig. S3**). Second, the threshold energy for applying boost potential to the ligand non-bonded interaction energy was set to the upper bound (i.e., *E* = *V*_min_+1/*k*). This enabled high enough boost potential to dissociate the ligand from the protein active site. In comparison, the bound ligand maintained the X-ray conformation during ∼200 ns testing simulations with the threshold energy set to the lower bound (i.e., *E* = *V*_max_). Moreover, higher acceleration was observed for the bound ligand as the input parameter *σ*_0*P*_ was increased from 1.0 kcal/mol to 6.0 kcal/mol and the ligand started to dissociate from the target site during the LiGaMD equilibration simulation with *σ*_0*P*_ = 4.0 kcal/mol (**Fig. S3**), which was thus used for production simulations. Third, a total of 10 ligand molecules (one in the X-ray bound conformation and another nine placed randomly in the solvent) were included in the system to facilitate ligand rebinding. This design was based on the fact that the ligand binding rate constant *k*_on_ is inversely proportional to the ligand concentration. The higher the ligand concentration, the faster the ligand binds, provided that the ligand concentration is still within its solubility limit. By applying the second boost potential to the unbound ligand molecules, protein and solvent, ligand rebinding could be sampled even during the LiGaMD equilibration simulations (**Figs. S3D** and **S3F**).

With the above settings, LiGaMD_Dual simulations were able to capture repetitive dissociation and binding of the benzamidine inhibitor in trypsin within 1 μs simulation time (**Figs. 5A, S4** and **S5** and **Movies S1-S5**). In five independent LiGaMD_Dual simulations as summarized in **Table 4**, the average of the GaMD boost potential *ΔV* was ∼21-22 kcal/mol with ∼4.2-4.3 kcal/mol standard deviation. The ligand dissociated for 3-11 times and rebound for 3-10 times during the five 1 μs LiGaMD_Dual simulations. In the X-ray bound conformation, the distance between the N atom in benzamidine and CG atom of Asp189 in trypsin was 3.9 Å. During the LiGaMD_Dual simulations, the bound ligand dissociated from the protein and diffused into the bulk solvent with the benzamidine:N – ASP189:CG distance increased up to ∼60 Å (**Figs. 5A** and **S5**), similarly for the ligand RMSD relative to the X-ray conformation (**Fig. S4**). Then after sufficient sampling of the bulk solvent space, one of the ten ligand molecules rebound to the protein with a salt bridge formed between the charged benzamidine and side chain of protein residue Asp189, for which the benzamidine:N – Asp189:CG distance and ligand RMSD dropped back to ∼3.9 Å (**Fig. 5A**) and ∼1.0 Å (**Fig. S4**), respectively. When the benzamidine:N – Asp189:CG distance dropped below a threshold value (3.7 Å defined here), atomic coordinates, velocities and forces of the bound ligand were swapped with those of the original bound ligand in the simulation starting structure (denoted “Lig0”). Upon dissociation of Lig0 from the protein, the high concentration of ten ligand molecules in the solvent facilitated ligand rebinding to the protein. Through such cycles, repetitive dissociation and binding of the benzamidine ligand in trypsin were efficiently sampled in the LiGaMD_Dual simulations.

**Table 4.** Summary of LiGaMD_Dual production simulations performed on the benzamidine ligand binding to trypsin. *ΔV* is the GaMD boost potential. *N*_*D*_ and *N*_*B*_ are the number of host-guest dissociation and binding events recorded from individual simulations.

| System | Method | ID | Length (ns) | $\Delta V$ (kcal/mol) | $N_D$ | $N_B$ |
| --- | --- | --- | --- | --- | --- | --- |
| Trypsin - benzamidine | LiGaMD_Dual | Sim1 | 1000 | $20.93 \pm 4.24$ | 3 | 3 |
| | | Sim2 | 1000 | $21.64 \pm 4.67$ | 5 | 4 |
| | | Sim3 | 1000 | $20.99 \pm 4.23$ | 11 | 10 |
| | | Sim4 | 1000 | $20.87 \pm 4.18$ | 4 | 4 |
| | | Sim5 | 1000 | $20.94 \pm 4.18$ | 4 | 4 |

**Fig. 5.**
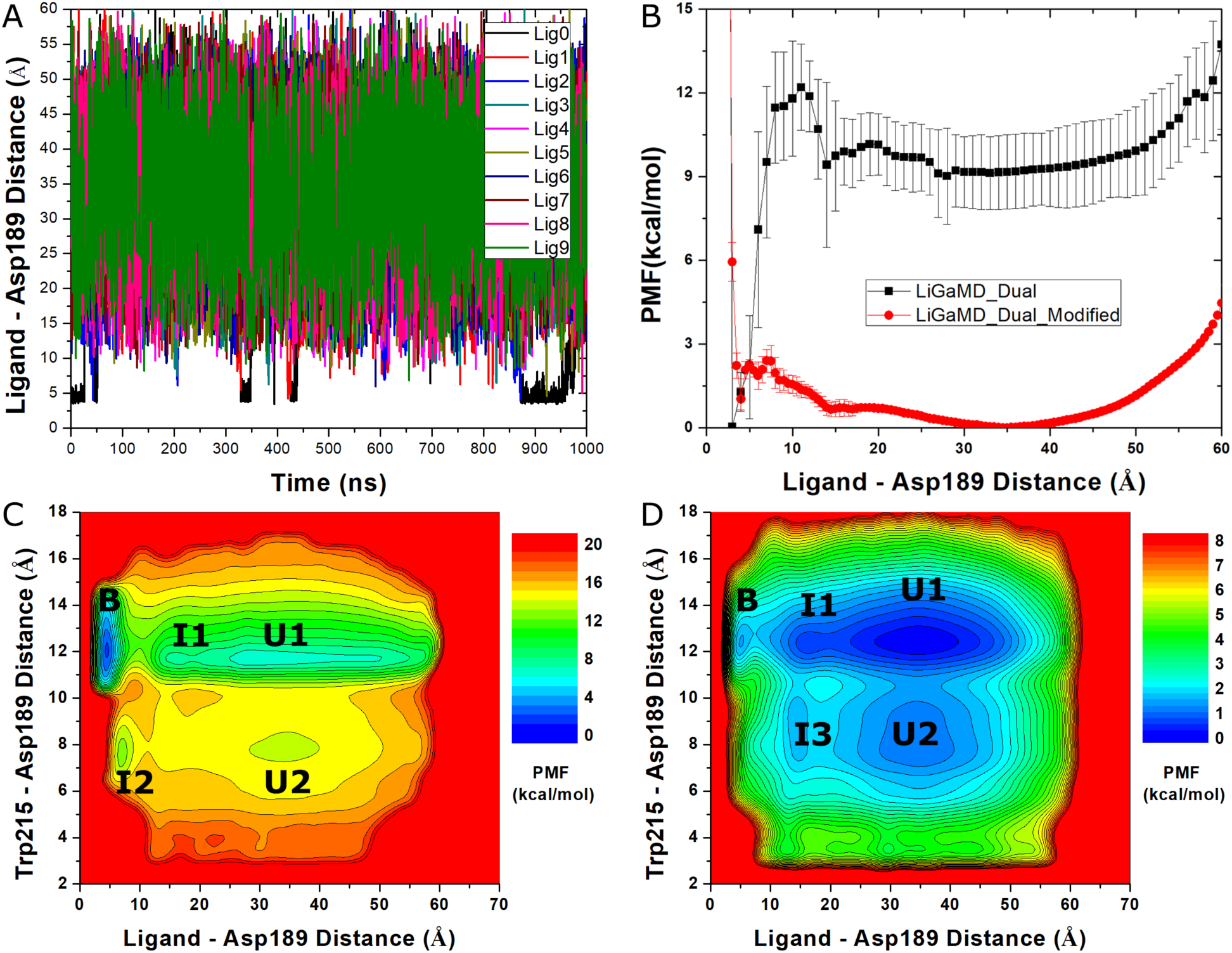
LiGaMD_Dual simulations have captured repetitive binding and unbinding of the benzamidine ligand in the trypsin protein: (A) time courses of distances between the N atom in benzamidine and CG atom of Asp189 in trypsin calculated from a representative 1 μs LiGaMD_Dual simulation (Sim2 in **Table 5**). Distance plots of the other simulations are provided in **Figure S5**. (B) The original (reweighted) and modified (no reweighting) PMF profiles of the distance between the N atom of benzamidine ligand and CG atom of protein residue Asp189. (C) Reweighted and (D) modified 2D PMF profiles of the benzamidine:N – Asp189:CG and Trp215:NE – Asp189:CG atom distances. The PMF profiles were calculated by combining all five 1000 ns LiGaMD_Dual simulations.

Next, we calculated PMF free energy profiles to characterize ligand binding to trypsin. Since plots of the benzamidine:N – Asp189:CG distance and ligand RMSD depicted the ligand binding processes similarly (**Figs. S4** and **S5**) and extra efforts were needed to calculate symmetry-corrected RMSD of benzamidine, the PMF profiles were calculated for the benzamidine:N – Asp189:CGdistance. The resulting reweighted and modified 1D PMF profiles were shown in **Fig. 5B**. In the reweighted PMF profile, three low-energy wells were identified for the ligand in the Bound, Intermediate and Unbound states, for which the benzamidine:N – Asp189:CG distance was centered around ∼4.5 Å, ∼15 Å and ∼35 Å, respectively. A high energy barrier of 12.17 ± 1.54 kcal/mol was observed for the ligand dissociation (**Table S5**). In the modified PMF profile without reweighting, the energy barrier decreased to 1.37 ± 0.56 kcal/mol for ligand dissociation and the energy wells became significantly shallower for the ligand in the Bound, Intermediate and Unbound states. This justified enhanced sampling of protein-ligand binding and unbinding in the LiGaMD_Dual simulations.

**Table 5.**
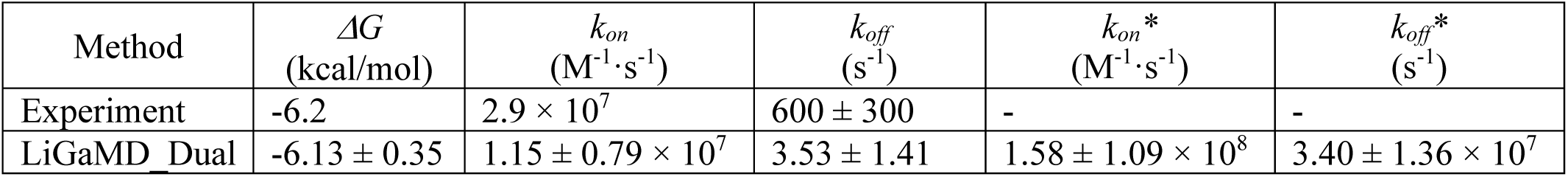
Comparison of trypsin protein-ligand binding free energy and kinetic rates obtained from experimental data and LiGaMD_Dual simulations. *ΔG* is the ligand binding free energy. *k*_*on*_ and *k*_*off*_ are the kinetic dissociation and binding rate constants, respectively, from experimental data or LiGaMD_Dual simulations with reweighting using Kramers’ rate theory. *k*_*on*_*\** and *k*_*off*_*\** are the accelerated kinetic dissociation and binding rate constants calculated directly from LiGaMD_Dual simulations without reweighting.

Furthermore, we computed 2D PMF profiles to analyze conformational changes of the protein upon ligand binding. Upon close examination of the system trajectories, residue Trp215 and its associated loop in trypsin underwent the largest conformational changes during ligand binding (**Movies S1-S5**). In this regard, a distance between the N_ε_ atom in the Trp215 side chain and CG atom of Asp189 was selected as another reaction coordinate. The calculated reweighted and modified 2D PMF profiles of the benzamidine:N – Asp189:CG and Trp215:NE – Asp189:CG atom distances are plotted in **Fig. 5C** and **5D**, respectively. Five 1000 ns LiGaMD_Dual simulations were combined for calculating the PMF profiles. The reweighted and modified 2D PMF profiles calculated from five individual LiGaMD_Dual simulations separately are also shown in **Figs. S6** and **S7**, respectively.

In the reweighted 2D PMF, low-energy wells were identified for the trypsin-benzamidine system in the Bound, Intermediate I1, Intermediate I2, Unbound U1 and Unbound U2 states, for which the (benzamidine:N – Asp189:CG, Trp215:NE – Asp189:CG) atom distances were centered around (4.0 Å, 12.5 Å), (17.0 Å, 12.5 Å), (7.5 Å, 7.5 Å), (35.0 Å, 12.5 Å) and (35.0 Å, 8.0 Å), respectively (**Fig. 5C**). In the modified 2D PMF profile (**Fig. 5D**), low-energy wells were also identified for the system in the Bound, Intermediate I1, Unbound U1 and Unbound U2 states in similar locations as in the reweighted 2D PMF. A distinct low-energy well was identified for the system in an Intermediate I3 state in the modified PMF with (14.0 Å, 9.0 Å) for the (benzamidine:N – Asp189:CG, Trp215:NE – Asp189:CG) atom distances, rather than the I2 state in the reweighted 2D PMF. These low-energy wells were also observed in the reweighed and modified PMF profiles of five individual LiGaMD_Dual simulations (**Figs. S6** and **S7**).

Next, we extracted six representative low-energy conformations of the trypsin-benzamidine system as identified from the PMF profiles, including the “Bound” **(Fig. 6A)**, “Intermediate I1” **(Fig. 6B)**, “Intermediate I2” **(Fig. 6C)**, “Intermediate I3” **(Fig. 6D)**, “Unbound U1” **(Fig. 6E)**, and “Unbound U2” **(Fig. 6F)**. In the Bound conformational state, the benzamidine ligand bound to protein residue Asp189 in the S1 pocket as in the X-ray crystal structure. Protein residue Trp215 closed the S1* pocket, but left the S1 pocket open, which was located between the Asp189 loop and the Trp215 loop (**Fig. 6A**). In the Intermediate I1 state, the benzamidine ligand bound to a site out of the Trp215 “gate”, being close to the His57-Asp102-Ser214 catalytic triad (**Fig. 5B**). In the Intermediate I3 state, residue Trp215 flipped its side chain to close the S1 pocket, while the S1* pocket became open with rearrangement of the Trp215 loop compared with the X-ray structure. The benzamidine ligand bound to the open S1* pocket (**Fig. 6C**). In the Intermediate I3 state, the side chain of protein residue Trp215 rotated to a conformation perpendicular to the X-ray conformation such that both S1 and S1* pockets became open and the benzamidine ligand bound to a site in close proximity of Trp215 and catalytic residues Asp102 and Ser214 (**Fig. 6D**). In the Unbound U1 state, trypsin adopted a conformation similar to the X-ray structure with the S1 pocket open and the S1* pocket closed by Trp215, while the benzamidine ligand was found far away from the protein (**Fig. 6E**). In the Unbound U2 state, trypsin flipped its side chain to close the S1 pocket, similar to the Intermediate I2 conformation, while the benzamidine ligand stayed far away from the protein (**Fig. 6F**).

**Fig. 6.**
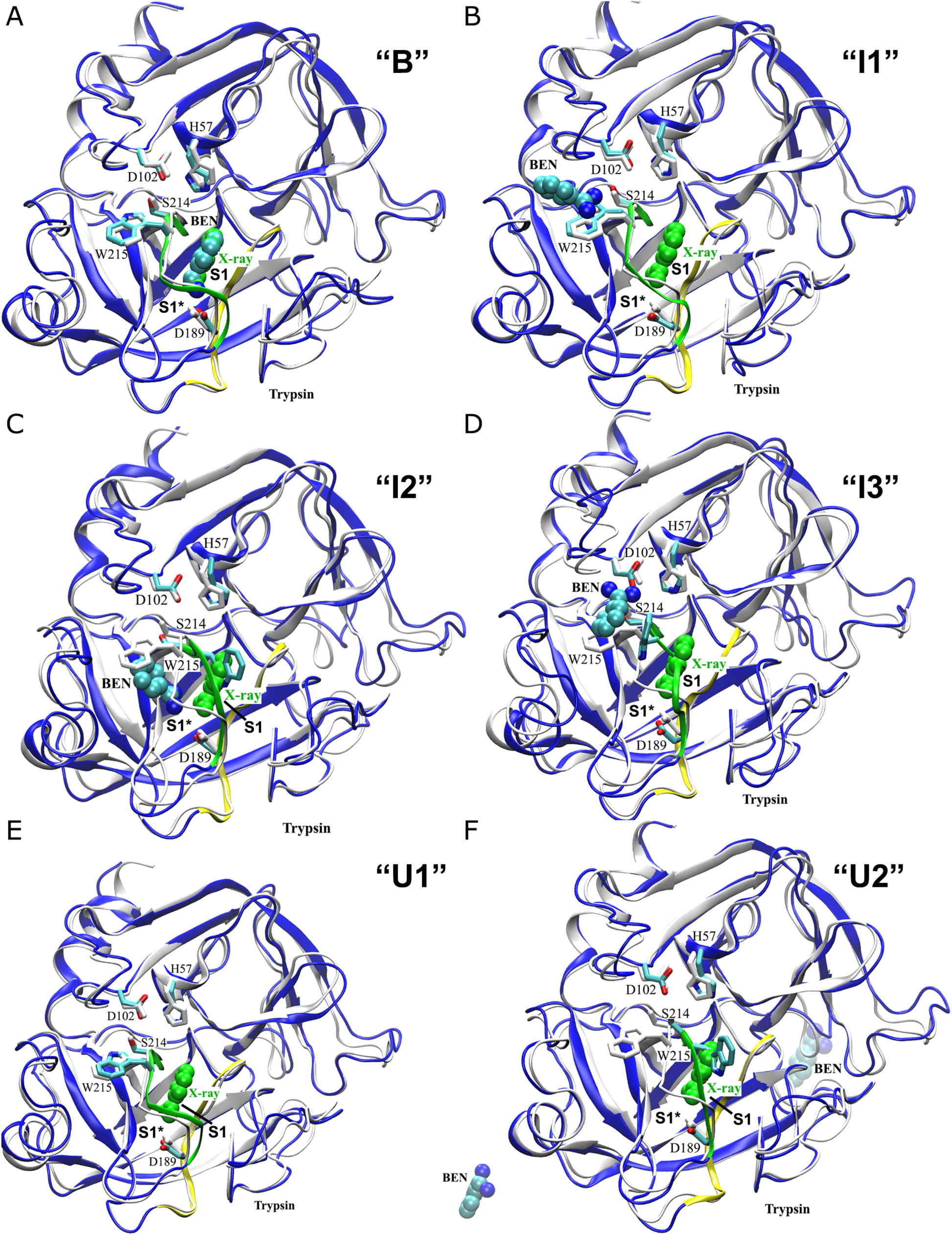
Six representative low-energy conformational states as identified from PMF profiles of benzamidine (BEN) ligand (spheres) binding to trypsin (blue ribbons): (A) “Bound (B)”, (B) “Intermediate 1 (I1)”, (C) “Intermediate 2 (I2)”, (D) “Intermediate 3 (I3)”, (E) “Unbound (U1)”, and (F) “Unbound (U2)”. Reference X-ray conformations of the ligand and protein (PDB: 3PTB) are shown in green spheres and grey ribbons, respectively. Protein residues Asp189 and Trp215 that are important for ligand binding and those in the catalytic triad (His57, Asp102 and Ser214) are highlighted in thick sticks.

In addition to the 1D and 2D PMF profiles, 3D PMF was calculated from each individual 1 μs LiGaMD_Dual simulation of benzamidine binding to trypsin in terms of displacements of the benzamidine N atom from the CG atom in protein residue Asp189 in the X, Y and Z directions. We then calculated the ligand binding free energy from each 3D reweighted PMF (see **Methods**). The average of resulting free energy values was -6.13 kcal/mol and the standard deviation was 0.35 kcal/mol. This was in excellent agreement with the experimental value of -6.2 kcal/mol for the ligand binding free energy of benzamidine in trypsin^58^ (**Table 5**).

### Pathways and kinetics of Ligand Binding to Trypsin

With accurate prediction of the ligand binding free energy, we analyzed the LiGaMD_Dual simulations further to determine the pathways and kinetic rate constants of benzamidine binding to trypsin. For each 1 μs simulation trajectory, structural clustering was performed on snapshots of the diffusing ligand molecules using the DBSCAN algorithm and the resulting structural clusters were reweighted to obtain energetically significant pathways of the ligand (see **Methods**). **Fig. 7** depicts the dissociation and binding pathways of the benzamidine ligand in trypsin obtained from the five individual LiGaMD_Dual simulations. The ligand clusters along each pathway were colored according to the reweighted PMF values in a blue (0 kcal/mol)-white (7.5 kcal/mol)-red (15.0 kcal/mol) scale. The lowest-energy ligand cluster was consistently identified at the target site with benzamidine forming ionic interaction with trypsin residue Asp189 as determined in the X-ray crystal structure (**Fig. 7**).

**Fig. 7.**
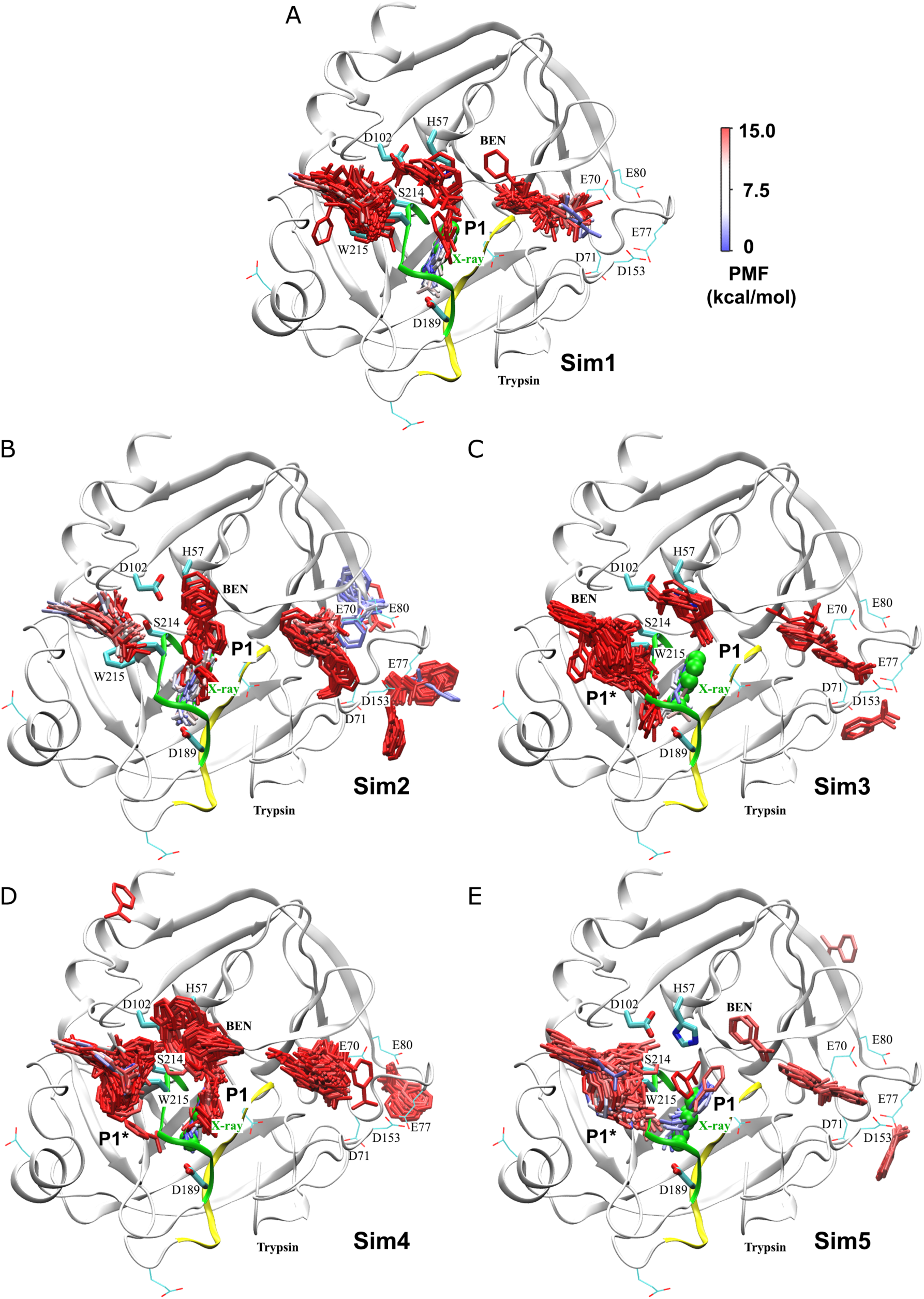
Pathways of the benzamidine (BEN) ligand (sticks) in trypsin (ribbons) obtained from structural clustering of five 1000 *ns* LiGaMD_Dual simulations: (A) Sim1, (B) Sim2, (C) Sim3, (D) Sim4 and (E) Sim5. The ligand is represented by sticks that are colored by reweighted PMF values of the ligand clusters in a blue (0 kcal/mol)-white (7.5 kcal/mol)-red (15.0 kcal/mol) scale. X-ray conformation of the ligand is shown in green spheres for reference. Protein residues Asp189 and Trp215 that are important for ligand binding and those in the catalytic triad (His57, Asp102 and Ser214) are highlighted in thick sticks. Other negatively charged residues in the protein including Glu70, Asp71, Glu77, Glu80 and Asp153 are shown in thin sticks. The protein loops containing residues Asp189 and Trp215 are highlighted in yellow and green, respectively.

During the LiGaMD_Dual simulations, benzamidine first dissociated predominantly through the protein opening between the Asp189 loop and the Trp215 loop, denoted pathway “P1” (**Fig. 7**). Then benzamidine molecules could rebind spontaneously to trypsin. Negatively charged residues on the protein surface, including Glu70, Asp71, Glu77, Glu80 and Asp153, appeared to steer the positively charged ligand towards the enzyme catalytic site formed by residues His57-Asp102-Ser214. This was consistent with previous findings of “electrostatic steering” in ligand recognition by proteins^59^. Subsequently, the ligand bound to the target site via two pathways, one being the pathway “P1” and the other connecting the S1* pocket and the Trp215 side chain gate as closed in the X-ray structure (denoted pathway “P1*”) (**Fig. 7**). In Sim1 and Sim2, because the Trp215 side chain maintained the X-ray conformation and closed the S1* pocket, the benzamidine ligand dissociated and bound to the S1 pocket repetitively through the P1 pathway (**Figs. 7A** and **7B** and **Movies S1** and **S2**). In the other three simulations Sim3-Sim5, the Trp215 side chain underwent conformational changes to open either the S1 or S1* pocket or both pockets. Benzamidine dissociated and bound to the protein via either of the P1 and P1* pathways (**Figs. 7C-7E** and **Movies S3-S5**). Notably, binding of the ligand was able to switch from pathway P1* to pathway P1 via the S1* binding pocket to the final target site in the S1 pocket as observed near 286 ns in the Sim5 trajectory (**Movie S5**). In addition, two ligand molecules were found to bind both the S1 and S1* pockets simultaneously during 310.5 ns-333.5 ns in the Sim5 trajectory (**Movie S5** and **Fig. S5E**).

To calculate kinetic rate constants of benzamidine binding to trypsin, we recorded the time periods for the ligand found in the bound (τ_B_) and unbound (τ_U_) states throughout the LiGaMD_Dual simulations (**Table S4**). Since a total of 10 ligand molecules and 10,478 water molecules were included in the simulation system, the ligand concentration was 0.053 M in the simulations. Without reweighting of the LiGaMD_Dual simulations, the benzamidine ligand binding (*k*_*on*_*\**) and dissociation (*k*_*off*_*\**) rate constants were calculated as 1.58 ± 1.09 × 10^8^ M×s^-1^ and 3.40 ± 1.36 × 10^7^ s^-1^, respectively.

Following the same protocol as described for analyzing the host-guest binding simulations, we reweighted the trypsin-benzamidine binding simulations to calculate acceleration factors of the ligand binding and dissociation (**Table S5**) and recover the original kinetic rate constants using the Kramers’ rate theory. The original (reweighted) and modified PMF profiles of protein-ligand distance are shown in **Fig. 5B**. The dissociation free energy barrier (Δ*F*_*off*_) decreased by ∼90% from 12.17 ± 1.54 kcal/mol in the reweighted PMF profile to 1.37 ± 0.56 kcal/mol in the modified PMF profile (**Table S5**). On the other hand, the free energy barrier for ligand binding (Δ*F*_*on*_) decreased from 3.04 ± 2.04 kcal/mol in the reweighted profile to 2.40 ± 0.41 kcal/mol in the modified PMF profile (**Fig. 5B** and **Table S5**). Furthermore, curvatures of the reweighed (*w*) and modified (*w*^∗^, no reweighting) free energy profiles were calculated near the guest bound (“B”) and unbound (“U”) low-energy wells and the energy barrier (“Br”), as well as the ratio of apparent diffusion coefficients calculated from the LiGaMD_Dual simulations with reweighting (*D*) and without reweighting (modified, *D*^∗^) (**Table S5**). According to the Kramers’ rate theory, the ligand binding and acceleration were accelerated by 13.76 and 9.62 × 10^7^ times, respectively. Therefore, the reweighted *k*_*on*_ and *k*_*off*_ were calculated as 1.15 ± 0.79 × 10^7^ M^-1^×s^-1^ and 3.53 ± 1.41 s^-1^, respectively. They were comparable to the experimental data^58^ of 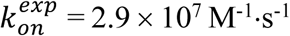 and 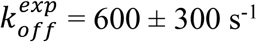 (**Table 5**).

## Discussion

A new LiGaMD method has been developed to selectively boost the ligand non-bonded interaction potential energy, which enables enhanced sampling simulations of repetitive ligand dissociation and binding as demonstrated on the host-guest and protein-ligand binding model systems. LiGaMD provides a promising approach for simultaneously calculating the free energy and kinetic rate constants of ligand binding.

For host-guest binding, LiGaMD and dual-boost LiGaMD (LiGaMD_Dual) significantly improved the sampling efficiency compared with the previous GaMD_Tot, GaMD_Dual, GaMD_NB and GaMD_NB_Dual algorithms (**Table 1**). For simulations that captured multiple events of ligand binding and dissociation, the accuracy of calculated ligand binding free energies and kinetic rate constants was similar to those obtained from much longer cMD simulations as compared with the experimental data. Notably, the CD host exhibited distinct structural dynamics when modeled with the different GAFF and q4MD force fields. Nevertheless, similar results were obtained from LiGaMD and cMD simulations provided the same force field. Therefore, LiGaMD only enhanced conformational sampling of the studied systems, but did not change statistics related to the force field. Apart from enhanced sampling, it remains critical to develop accurate force fields for systems of our interest such as the proteins and ligand molecules.

For the trypsin model protein-ligand binding system, the threshold energy for applying LiGaMD boost potential to non-bonded potential energy of the bound ligand was set to the upper bound so that high enough acceleration was obtained to allow for ligand dissociation. Multiple ligand molecules (e.g., 10 in the present simulations) were included in the system to facilitate binding. This was based on the fact that higher ligand concentration would lead to faster binding rate constant *k*_on_. Multiple events of ligand dissociation and binding were observed in each of the five independent 1 μs LiGaMD_Dual simulations of the trypsin-benzamidine system. Trypsin residue Trp215 appeared to be a gate for ligand binding through the P1 and P1* pathways.

The low-energy conformational states (**Fig. 6**) and ligand pathways (**Fig. 7**) of the trypsin-benzamidine system identified from LiGaMD_Dual simulations were mostly consistent with those obtained from previous simulation studies, especially the MSM^22^. However, LiGaMD_Dual simulations further revealed two novel findings. First, two benzamidine molecules were able to bind both the S1 and S1* pockets simultaneously during one of the five LiGaMD_Dual simulation trajectories (**Movie S5**). While this finding could result from a relatively high ligand concentration (0.053 M) with 10 benzamidine molecules in the simulation system, such rare event is worthy further investigation in the future. Second, binding of the benzamidine ligand was able to switch from the P1* pathway to the P1 pathway in the LiGaMD_Dual simulations. The ligand bound to the S1 pocket via an intermediate site in the S1* pocket. LiGaMD_Dual appeared to provide improved sampling compared with MD simulations used in the MSM so that such binding process could be captured. This LiGaMD_Dual simulation finding also suggested that the final target binding site of benzamidine is located in the S1 pocket as determined in the X-ray crystal structure. Ligand binding in the X-ray conformational state corresponded the global free energy minimum in the calculated PMF profiles.

Furthermore, the ligand binding free energy calculated from five 1 μs LiGaMD_Dual simulations Δ*G*^4^ = -6.13 ± 0.35 kcal/mol was in excellent agreement with the experimental value 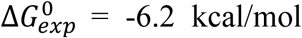. The ligand kinetic rate constants calculated by reweighting of the LiGaMD_Dual simulations were *k*_*on*_ = 1.15 ± 0.79 × 10^7^ M^-1^×s^-1^ and *k*_*off*_ = 3.53 ± 1.41 s^-1^. The binding rate constant *k*_on_ agreed excellently with the experimental value 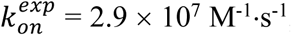, while the dissociation rate constant *k*_off_ appeared to be slower than the experimental value 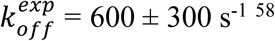 (**Table 5**).

In comparison, the ligand dissociation rate constant estimated from five 1 μs LiGaMD_Dual simulations *k*_*off*_ = 3.53 ± 1.41 s^-1^ was similar to the value of *k*_off_ = 9.1 ± 2.5 s^-1^ obtained from previous metadynamics simulations also performed at a total of 5 μs length^16a^. However, the benzamidine binding free energy calculated from LiGaMD_Dual simulations Δ*G*^0^ = -6.13 ± 0.35 kcal/mol was more accurate than the values of 8.5 ± 0.7 kcal/mol obtained from separate funnel metadynamics simulations^8^ and -5.06 kcal/mol from the SITSMD simulations^24^, as compared with experimental data 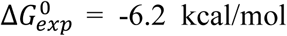. In this regard, previous MSM predicted accurate ligand binding free energy Δ*G*^0^ = -6.05 ± 1 kcal/mol and binding rate constant *k*_on_ = 6.4 ± 1.6 × 10^7^ M^-1^·s^-1^, while the model predicted *k*_off_ = 131 ± 109 × 10^2^ s^-1^ appeared to be faster than the experimental value^22^. Both MSM and LiGaMD_Dual could be applied to calculate the ligand binding free energy and kinetic rate constants simultaneously, but more expensive and significantly longer simulations were needed for the MSM (∼150 μs in total) than for the LiGaMD_Dual simulations (1 μs × 5).

In summary, LiGaMD provides a promising approach to calculate both free energy and kinetic rate constants of ligand binding simultaneously. The ligand binding free energy is calculated based on unconstrained enhanced sampling and 3D PMF profiles, being distinct from previous methods such as the TI^3^, FEP^4^ and funnel metadynamics^8^. Beyond thermodynamics, LiGaMD also presents a new approach to estimate the ligand kinetic dissociation and binding rate constants. While LiGaMD has been demonstrated on host-guest and trypsin protein-ligand binding model systems, the method awaits further testing on many other ligand binding systems, notably membrane proteins. Although a total number of 10 ligand molecules has been included in the trypsin-benzamidine simulations to facilitate ligand binding, it will be more systematic to optimize the number of ligand molecules for future simulations according to the ligand solubility. Furthermore, a distance cutoff has been implemented in the current LiGaMD to determine when a ligand molecule binds to the protein target site and higher boost potential will be applied accordingly to enable ligand dissociation. More metrics that can serve in this purpose for a wide range of ligand binding systems with different chemical and physical properties will be implemented in the future. These developments are expected to further improve the LiGaMD method for applications in characterizing both ligand binding thermodynamics and kinetics, which should facilitate computer-aided drug design.

## Appendix A Gaussian accelerated molecular dynamics (GaMD)

Consider a system with *N* atoms at positions 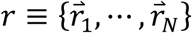. When the system potential *V*(*r*) is lower than a reference energy *E*, the modified potential *V*^∗^(*r*) of the system is calculated as:

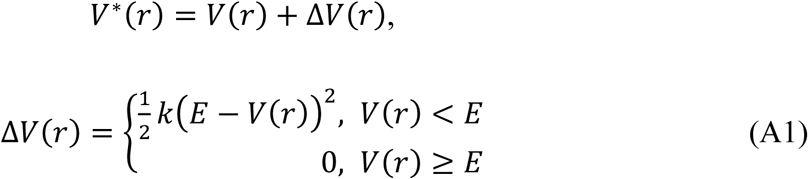

where *k* is the harmonic force constant. The two adjustable parameters *E* and *k* are automatically determined based on three enhanced sampling principles^25^. The reference energy needs to be set in the following range:

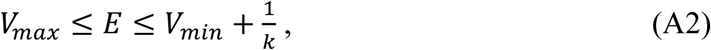

where *V*_*max*_ and *V*_*min*_ are the system minimum and maximum potential energies. To ensure that Eqn. (A2) is valid, *k* has to satisfy: 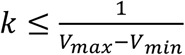 Let us define 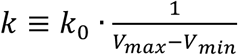, then 0 < *k*_0_ ≤ 1. The standard deviation of Δ*V* needs to be small enough (i.e., narrow distribution) to ensure proper energetic reweighting^60^: *σ*_Δ*V*_ = *k*(*E* − *V*_*avg*_)*σ*_*V*_ ≤ *σ*_0_ where *V*_*avg*_ and *σ*_*V*_ are the average and standard deviation of the system potential energies, *σ*_Δ*V*_ is the standard deviation of Δ*V* with *σ*_0_ as a user-specified upper limit (e.g., 10*k*_*B*_T) for proper reweighting. When *E* is set to the lower bound *E=V*_*max*_, *k*_0_ can be calculated as:

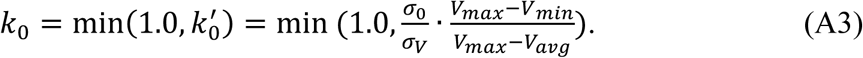

Alternatively, when the threshold energy *E* is set to its upper bound 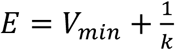, *k*_0_ is set to:

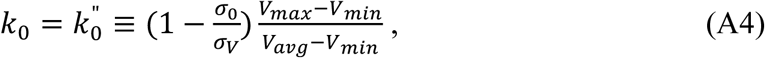

if 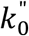 is found to be between *0* and *1*. Otherwise, *k*_0_ is calculated using Eqn. (A3).

GaMD provides options to boost only the total potential boost (GaMD_Tot), only the dihedral potential energy (GaMD_Dih), both the total and dihedral potential energies (GaMD_Dual), the non-bonded potential energy (GaMD_NB), both the non-bonded potential and dihedral energies (GaMD_Dual_NB), only non-bonded potential energy of the bound ligand (LiGaMD), and both the ligand non-bonded potential energy of the bound ligand and the remaining potential energy of the entire system (LiGaMD_Dual). The dual-boost simulation generally provides higher acceleration than the other single-boost simulations for enhanced sampling. The simulation parameters comprise of settings for calculating the threshold energy values and the effective harmonic force constants of the boost potentials.

## Appendix B Energetic reweighting of GaMD simulations

For energetic reweighting of GaMD simulations to calculate potential of mean force (PMF), the probability distribution along a reaction coordinate is written as *p*^∗^(*A*). Given the boost potential Δ*V*(*r*) of each frame, *p*^∗^(*A*) can be reweighted to recover the canonical ensemble distribution, *p*(*A*), as:

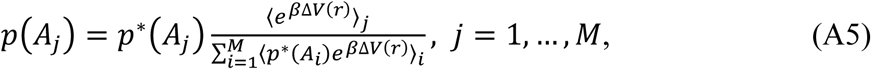

where *M* is the number of bins, *β* = *k*_*B*_*T* and ⟨*e*^*β*Δ7*(r)*^⟩_*j*_ is the ensemble-averaged Boltzmann factor of Δ*V*(*r*) for simulation frames found in the *j*^th^ bin. The ensemble-averaged reweighting factor can be approximated using cumulant expansion:

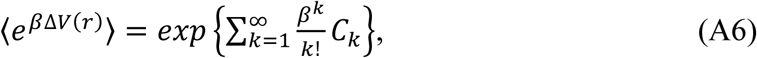

where the first two cumulants are given by:

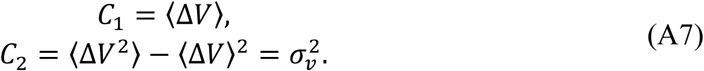

The boost potential obtained from GaMD simulations usually follows near-Gaussian distribution^33a^. Cumulant expansion to the second order thus provides a good approximation for computing the reweighting factor^25, 60^. The reweighted free energy *F*(*A*) = −*k*_*B*_*T* ln *p*(*A*) is calculated as:

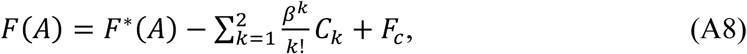

where *F*^∗^(*A*) = −*k*_*B*_*T* ln *p*^∗^(*A*) is the modified free energy obtained from GaMD simulation and *F*_/_ is a constant.

## Appendix C Reweighting of Biomolecular Kinetics with Kramers’ Rate Theory

A brief summary is provided here for reweighting of biomolecular kinetics from GaMD simulations with Kramers rate theory as described recently^26^. For a particle climbing over potential energy barriers, Kramers showed that the reaction rate depends on temperature and viscosity of the host medium^61^. The reaction rates were derived for both limiting cases of small and large viscosity. In the context of biomolecular simulations in aqueous medium, it is relevant for us to focus on the large viscosity limiting case. Biomolecules move in the high friction (“overdamping”) regime and energy barriers are much greater than *k*_*B*_*T* (*k*_*B*_ is the Boltzmann’s constant and *T* is temperature). In this case, the reaction rate is calculated as:

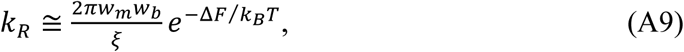

where *w*_*m*_ and *w*_*b*_ are frequencies of the approximated harmonic oscillators (also referred to as curvatures of free energy surface^45^) near the energy minimum and barrier, respectively, *ξ* is the apparent friction coefficient and Δ*F* is the free energy barrier of transition.

Without the loss of generality, we consider a 1D potential of mean force (PMF) free energy profile of a reaction coordinate *F*(*A*). Near minimum at *A*_*m*_, the free energy can be approximated by a harmonic oscillator^61^ of frequency *w*_*m*_, i.e., 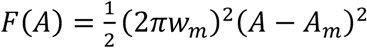. Near barrier at *A*_*b*_, the free energy is approximated as 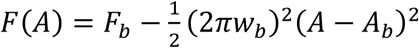, where *F*_*b*_ is the free energy at *A*_*b*_ and *w*_*b*_ is the frequency of the approximated harmonic oscillator. Then we can calculate *w*_*m*_ and *w*_*b*_ as:

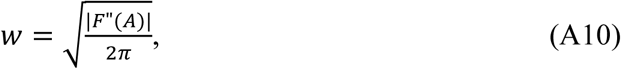

where *F*”(*A*) is the second-order derivative of the PMF profile.

The apparent friction coefficient *ξ* or diffusion coefficient *D* with *ξ* = *k*_*B*_*T*/*D* can be estimated as follows. First, we calculate a survival function *S*(*t*) as the probability that the system remains in an energy well longer than time *t*. In a direct approach^46^, we count the events that the system visits the energy well throughout a simulation. We record and measure the time intervals of each visiting event until the system escapes over an energy barrier. Then we have a time series *T*_*i*_, where *i*=1, 2, …, *N*, and *N* is the total number barrier transitions observed in the simulation. The time series is subsequently ordered such that 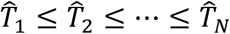. With that, the survival function is estimated as 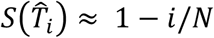, which is the probability that the system is trapped in the energy well for time longer than 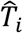. Alternatively, we can numerically calculate the time-dependent probability density of reaction coordinate *A, ρ*(*A, t*) by solving the Smoluchowski equation along 1D PMF profile of the reaction coordinate:

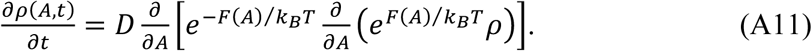

Then the survival function is calculated as 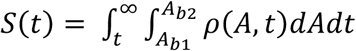, where *A*_*b*1_ and *A*_*b*2_ are two boundaries of the energy well. The initial condition is often set as the Boltzmann distribution of reaction coordinate *A* in the energy well, i.e., 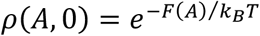.

Second, using the above survival functions, we estimate the effective kinetic rates as the negative of the slopes in linear fitting of the ln[*S*(*t*)] versus *t*, i.e., *k* = −*d*ln[*S*(*t*)]/*dt*. This is based on the assumption that the survival function exhibits exponential decay as observed in earlier studies^46, 62^. Finally, the apparent diffusion coefficient *D* is obtained by dividing the kinetic rate calculated directly using the transition time series collected from the simulation by that using the probability density solution of the Smoluchowski equation^46^.

The curvatures and energy barriers of the reweighted and modified free energy profiles, as well as the apparent diffusion coefficients, are calculated and used in Kramers’ rate equation to determine accelerations of biomolecular kinetics in the GaMD simulations^26^.

## Appendix D: Implementation of ligand Gaussian accelerated molecular dynamics

Ligand Gaussian accelerated molecular dynamics (LiGaMD) is currently implemented in the GPU version of AMBER 20^63^, but should be transferable to other molecular dynamics programs as well. LiGaMD provides enhanced sampling of protein-ligand binding and unbinding. Following is a list of the input parameters for a LiGaMD simulation:

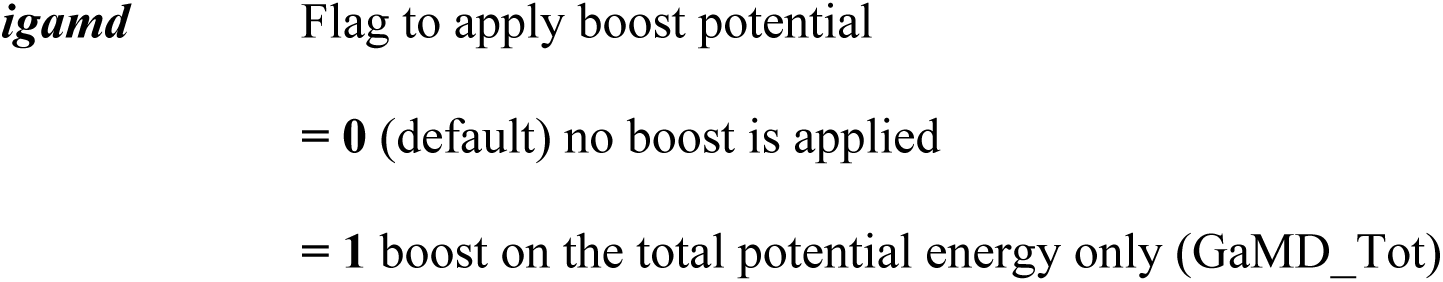
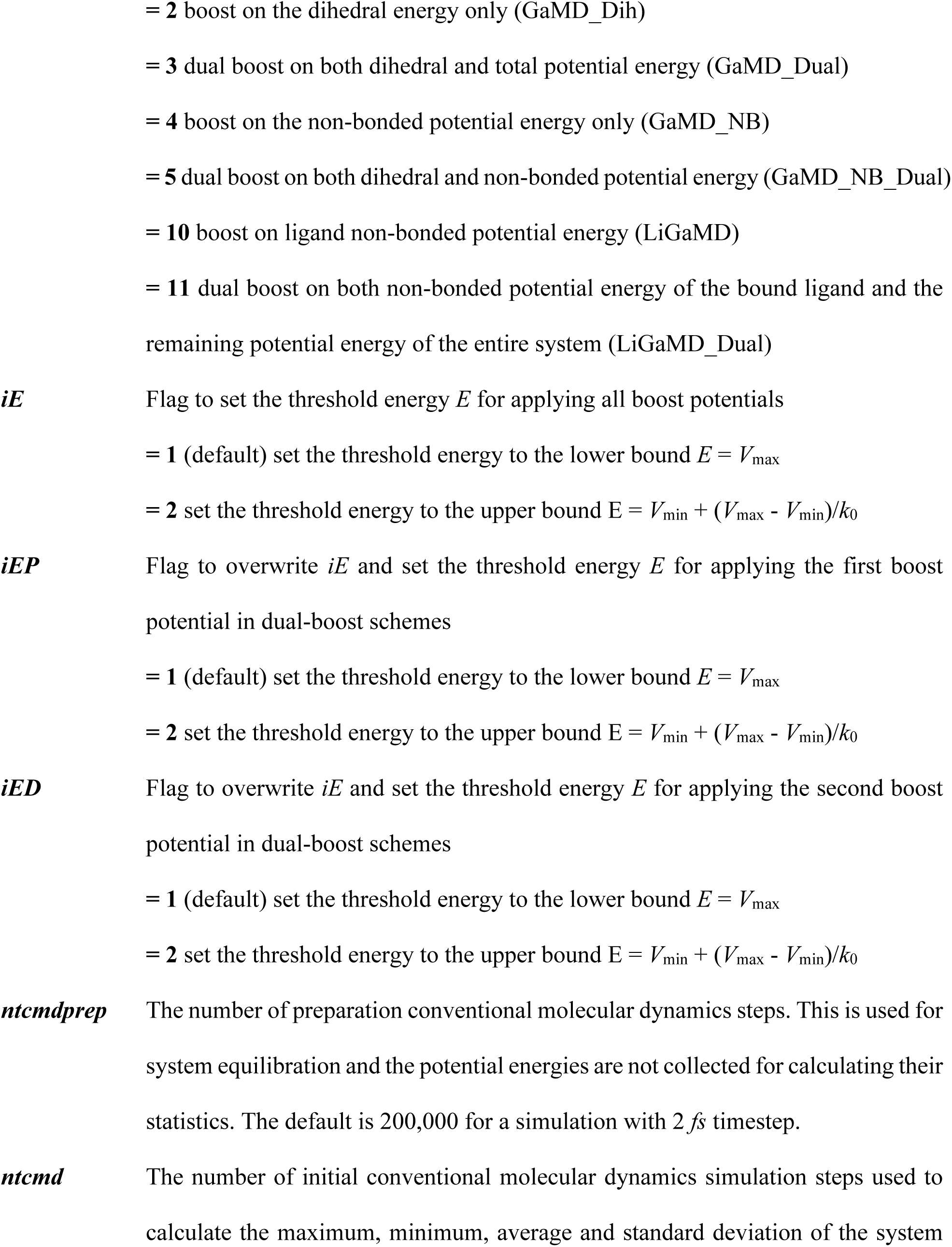
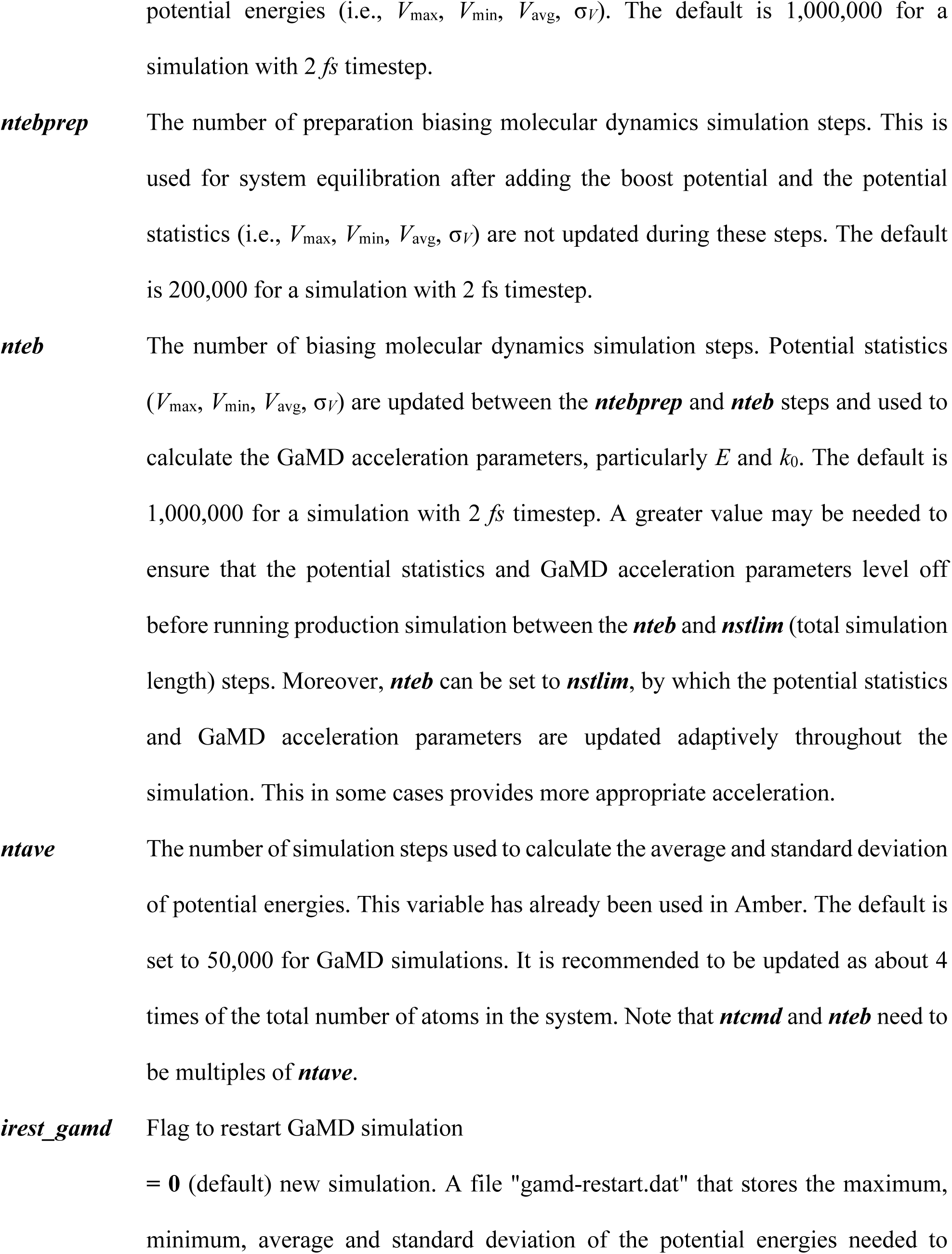
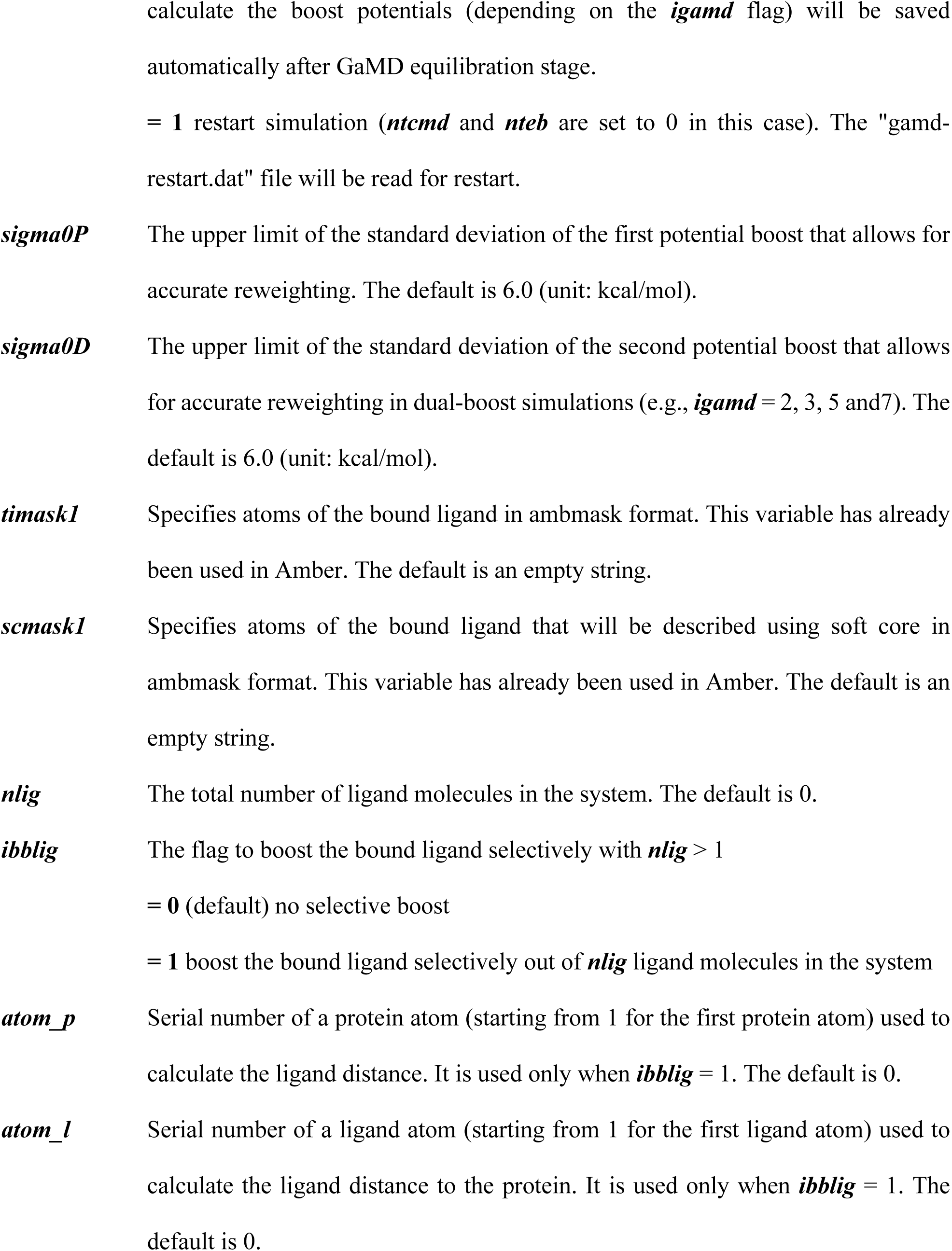
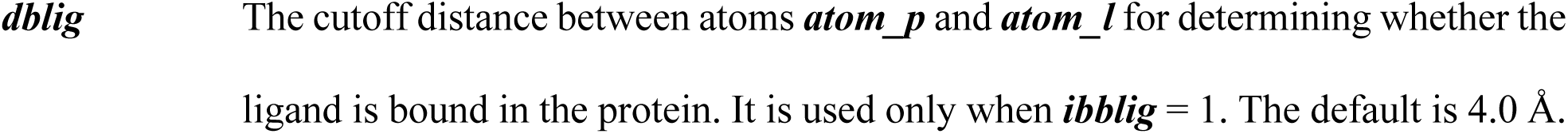

Example input parameters used in LiGaMD_Dual simulations of ligand binding to trypsin include the following:

~~~
  igamd = 11, irest_gamd = 0,
  ntcmd = 700000, nteb = 27300000, ntave = 140000,
  ntcmdprep = 280000, ntebprep = 280000,
  sigma0P = 4.0, sigma0D = 6.0, iEP = 2, iED=1,
  icfe = 1, ifsc = 1, gti_cpu_output = 0, gti_add_sc = 1,
  timask1 = ‘:225’, scmask1 = ‘:225’,
  timask2 = ‘‘, scmask2 = ‘‘,
  ibblig = 1, nlig = 10, atom_p = 2472, atom_l = 4, dblig = 3.7
~~~

The LiGaMD algorithm is summarized as the following:

~~~
LiGaMD {
  If (irest_gamd == 0) then
    For i = 1, …, ntcmd // run initial conventional molecular dynamics
      If (i >= ntcmdprep) Update Vmax, Vmin
      If (i >= ntcmdprep && i%ntave ==0) Update Vavg, sigmaV
    End
    Save Vmax,Vmin,Vavg,sigmaV to “gamd_restart.dat” file
    Calc_E_k0(iE,sigma0,Vmax,Vmin,Vavg,sigmaV)
    For i = ntcmd+1, …, ntcmd+nteb // Run biasing molecular dynamics simulation steps
      deltaV = 0.5*k0*(E-V)**2/(Vmax-Vmin)
      V = V + deltaV
      If (i >= ntcmd+ntebprep) Update Vmax, Vmin
      If (i >= ntcmd+ntebprep && i%ntave ==0) Update Vavg, sigmaV
      Calc_E_k0(iE,sigma0,Vmax,Vmin,Vavg,sigmaV)
    End
    Save Vmax,Vmin,Vavg,sigmaV to “gamd_restart.dat” file
  else if (irest_gamd == 1) then
    Read Vmax,Vmin,Vavg, sigmaV from “gamd_restart.dat” file
  End if
  lig0=1 // ID of the bound ligand
  For i = ntcmd+nteb+1, …, nstlim // run production simulation
    If (ibblig>0 && i%ntave ==0) then // swap the bound ligand with lig0 for selective boost
     For ilig = 1, …, nlig
       dlig = distance(atom_p, atom_l)
       If (dlig <= dblig) blig=ilig
     End
     If (blig != lig0) Swap atomic coordinates, forces and velocities of ligands *blig* with lig0
   End if
   deltaV = 0.5*k0*(E-V)**2/(Vmax-Vmin)
   V = V + deltaV
  End
}
Subroutine Calc_E_k0(iE,sigma0,Vmax,Vmin,Vavg,sigmaV) {
if iE = 1 :
    E = Vmax
    k0’ = (sigma0/sigmaV) * (Vmax-Vmin)/(Vmax-Vavg)
    k0 = min(1.0, k0’)
else if iE = 2 :
    k0” = (1-sigma0/sigmaV) * (Vmax-Vmin)/(Vavg-Vmin)
    if 0 < k0” <= 1 :
        k0 = k0”
        E = Vmin + (Vmax-Vmin)/k0
    else
        E = Vmax
        k0’ = (sigma0/sigmaV) * (Vmax-Vmin)/(Vmax-Vavg)
        k0 = min(1.0, k0’)
    end
end
}
~~~

## Acknowledgements

We appreciate the help of Prof. David Case for accessing the AMBER git repository to develop our new simulation algorithms. We thank Prof. Chia-en Chang and Dr. Zhiye Tang for kindly sharing molecular dynamics simulation files of the host-guest binding and valuable discussions. We thank Dr. Ferran Feixas for preliminary simulations and valuable discussions on the trypsin-benzamidine system. We also thank Prof. Darrin York and Dr. Taisung Lee for valuable discussions on coding in AMBER. This work used supercomputing resources with allocation award TG-MCB180049 through the Extreme Science and Engineering Discovery Environment (XSEDE), which is supported by National Science Foundation grant number ACI-1548562, and project M2874 through the National Energy Research Scientific Computing Center (NERSC), which is a U.S. Department of Energy Office of Science User Facility operated under Contract No. DE-AC02-05CH11231, and the Research Computing Cluster at the University of Kansas. This work was supported in part by the American Heart Association (Award 17SDG33370094), the National Institutes of Health (R01GM132572) and the startup funding in the College of Liberal Arts and Sciences at the University of Kansas.

## Supporting Information

Five supplementary **Tables S1 – S5**, seven **Figures S1** - **S7** and five **Movies S1 – S5** are provided in the supporting information. This information is available free of charge via the Internet at http://pubs.acs.org.

## TOC graphic

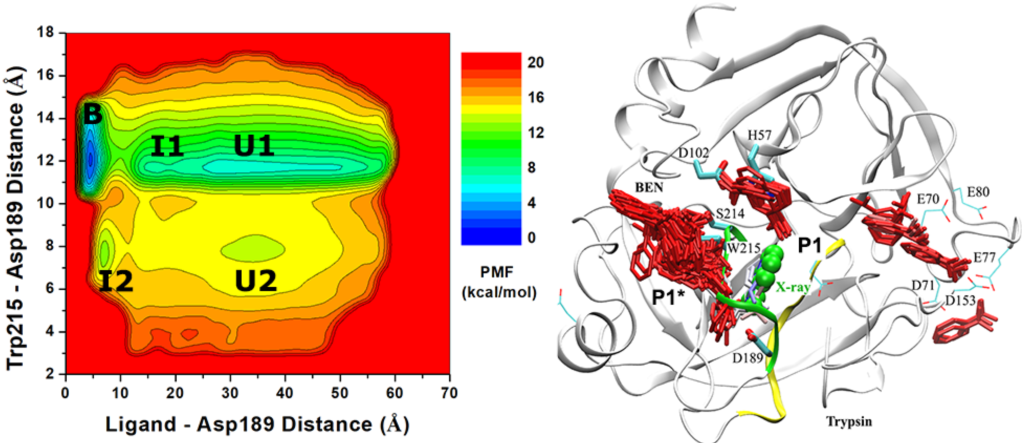

All-atom ligand Gaussian accelerated molecular dynamics (LiGaMD) simulations captured repeptitive binding of the benzamidine (BEN) ligand to the trypsin model protein, which enabled us to characterize the ligand binding free energy profiles, pathways and kinetic rate constants.

