## Supporting Information for "Ligand Gaussian accelerated molecular dynamics (LiGaMD): Characterization of ligand binding thermodynamics and kinetics"

**Table S1** Summary of host-guest binding thermodynamics and kinetics obtained from (A) experimental data and (B) cMD simulations from Ref. [1]. Results are obtained for the binding of aspirin and 1-butanol guests to the β-cyclodextrin (CD) host. In simulations, the CD host has been modeled with both the GAFF and q4MD force fields. $\Delta G$ is the ligand binding free energy. The $\Delta G_{comp1}$ and $\Delta G_{comp2}$ are ligand binding free energies calculated using two different algorithms as adapted from Ref. [1]. $k_{on}$ and $k_{off}$ are the kinetic dissociation and binding rate constants, respectively, with $N_D$ and $N_B$ being the number of host-guest dissociation and binding events collected from individual simulations.

| Host | Ligand | $\Delta G$ (kcal/mol) | $k_{on}$ ($\times 10^8$ M·s$^{-1}$) | $k_{off}$ ($\times 10^6$ s$^{-1}$) |
|---|---|---|---|---|
| CD | Aspirin | -3.74 ± 0.00 | 7.2 ± 0.04 | 1.3 ± 0.03 |
| | 1-Butanol | -1.67 ± 0.19 | 2.8 ± 0.8 | 38 ± 6 |

(A) Experimental data

| Host | Ligand | cMD | $N_D$ | $N_B$ | $\Delta G_{comp1}$ (kcal/mol) | $\Delta G_{comp2}$ (kcal/mol) | $k_{on}$ ($\times 10^8$ M·s$^{-1}$) | $k_{off}$ ($\times 10^6$ s$^{-1}$) |
|---|---|---|---|---|---|---|---|---|
| CD: GAFF | Aspirin | 9500 ns | 133 | 133 | -3.84 ± 0.35 | -2.27 ± 0.06 | 11 ± 0.1 | 24 ± 3 |
| | 1-Butanol | 6500 ns | 42 | 42 | 1.01 ± 0.59 | -0.41 ± 0.10 | 2.2 ± 0.1 | 110 ± 20 |
| CD: q4MD | Aspirin | 6000 ns | 17 | 18 | -6.33 ± 0.37 | -4.11 ± 0.05 | 32 ± 3 | 3.1 ± 0.9 |
| | 1-Butanol | 5000 ns | 89 | 89 | -1.33 ± 0.34 | -2.27 ± 0.02 | 15 ± 0.3 | 33 ± 0.7 |

(B) cMD simulations.

**Table S2** The guest binding and unbinding time periods ($\tau_B$ and $\tau_U$) recorded from LiGaMD simulations of host-guest binding systems.

| Host | Ligand | GaMD (300 ns x 3) | $\tau_B$ (ns) | $\tau_U$ (ns) |
|---|---|---|---|---|
| CD: GAFF | Aspirin | LiGaMD | 32.0228, 8.5966, 3.1112, 39.9596, 21.2604 | 9.9081, 194.0138, 86.3999 |
| | | LiGaMD_Dual | 8.2607, 0.796, 7.9322, 0.5776, 4.5163, 1.758, 5.9326 | 201.0795, 13.8457, 24.4303, 239.0854, 8.3765 |
| CD: q4MD | Aspirin | LiGaMD | 31.9081, 61.633, 43.8801, 8.2827, 16.9167, 20.3661, 14.1932, 19.3123, 27.4969 | 1.0855, 2.1165, 86.4148, 35.6304, 17.3932, 115.365, 89.763 |
| | | LiGaMD_Dual | 3.907, 30.0912, 36.3697, 1.4833, 10.1038, 1.7181, 12.8671, 56.2154, 3.9519, 8.4395, 8.9309, 4.4481, 4.2826, 26.9633, 38.3872, 20.3627, 9.6437, 44.3214 | 8.4001, 29.2999, 6.2442, 42.1422, 12.1334, 39.4287, 54.0979, 62.0065, 15.8736, 45.7133, 6.6287, 29.8733, 13.1733, 60.1833, 65.7046, 9.4815 |
| | 1-Butanol | LiGaMD | 5.4329, 2.5518, 11.281, 3.0967, 6.9, 2.606, 0.7952, 3.3571, 1.0371, 1.1861, 5.05, 0.9876, 1.275, 0.7967, 0.4231, 1.0814, 1.0247, 0.9549, 0.7596, 4.4845 | 5.1734, 72.0719, 51.3295, 15.962, 56.6445, 0.9302, 16.088, 12.9821, 19.5307, 84.7936, 62.3141, 27.7493, 39.8894, 61.5153, 32.1647, 24.3296, 24.2369, 38.1307, 27.2135, 79.995 |
| | | LiGaMD_Dual | 1.4312, 2.3294, 4.6188, 1.5447, 1.2259, 2.1735, 1.6123, 1.8385, 0.8007, 5.3053, 0.6236, 0.9946, 0.924, 0.3333, 4.6931, 1.8042, 0.8413, 2.2369, 1.7726, 2.692, 4.133, 1.7794, 8.4176, 2.262, 1.5251, 6.0076, 1.262, 1.5508, 1.1358, 1.6418 | 1.8388, 7.8776, 20.9217, 79.5963, 41.6825, 50.1632, 22.8479, 1.5881, 11.4997, 18.477, 65.5491, 16.961, 47.0128, 8.5518, 2.548, 2.8552, 5.14, 16.607, 5.6482, 41.906, 35.6489, 4.0918, 18.9406, 20.069, 14.6097, 8.0379, 85.2561, 4.8654, 24.6768, 45.8237 |

**Table S3** Energy barriers of host-guest dissociation ("off") and binding ("on") calculated from the reweighed (**Δ*F***) and modified (no reweighting, **Δ*F****) free energy profiles, curvatures of the reweighed (*w*) and modified ($w^*$) free energy profiles near the guest Bound ("B"), Barrier ("Br") and Unbound ("U") states, and the ratio of apparent diffusion coefficients calculated from the LiGaMD and LiGaMD_Dual simulations without reweighting (modified, $D^*$) and with reweighting (*D*). Results are listed for the following systems: (A) aspirin binding to CD with the GAFF force field, (B) aspirin binding to CD with the q4MD force field and (C) 1-butanol binding to CD with the q4MD force field.

| Sim | ΔF (kcal/mol) | | ΔF* (kcal/mol) | | w | | | w* | | | D*/D | |
|---|---|---|---|---|---|---|---|---|---|---|---|---|
| | Off | On | Off | On | B | Br | U | B | Br | U | Off | On |
| LiGaMD | 3.09 ± 0.37 | 0.61 ± 0.37 | 1.49 ± 0.35 | 1.87 ± 0.28 | 0.38 ± 0.21 | 0.08 ± 0.04 | 0.11 ± 0.01 | 0.32 ± 0.08 | 0.07 ± 0.08 | 0.15 ± 0.01 | 1.14 | 1.53 |
| LiGaMD_Dual | 2.06 ± 0.28 | 0.40 ± 0.34 | 0.59 ± 0.37 | 1.83 ± 0.28 | 0.49 ± 0.08 | 0.07 ± 0.05 | 0.12 ± 0.03 | 0.35 ± 0.03 | 0.05 ± 0.15 | 0.13 ± 0.01 | 1.23 | 0.85 |

(A) CD (GAFF) – Aspirin

| Sim | ΔF (kcal/mol) | | ΔF* (kcal/mol) | | w | | | w* | | | D*/D | |
|---|---|---|---|---|---|---|---|---|---|---|---|---|
| | Off | On | Off | On | B | Br | U | B | Br | U | Off | On |
| LiGaMD | 4.00 ± 0.34 | 0.97 ± 0.35 | 2.18 ± 0.28 | 1.95 ± 0.37 | 0.41 ± 0.08 | 0.09 ± 0.08 | 0.13 ± 0.03 | 0.32 ± 0.05 | 0.08 ± 0.02 | 0.13 ± 0.02 | 0.84 | 0.12 |
| LiGaMD_Dual | 4.46 ± 0.22 | 0.67 ± 0.39 | 2.25 ± 0.23 | 1.67 ± 0.24 | 0.46 ± 0.03 | 0.10 ± 0.12 | 0.18 ± 0.02 | 0.33 ± 0.02 | 0.08 ± 0.12 | 0.15 ± 0.00 | 0.92 | 1.80 |

(B) CD (q4MD) – Aspirin

| Sim | ΔF (kcal/mol) | | ΔF* (kcal/mol) | | w | | | w* | | | D*/D | |
|---|---|---|---|---|---|---|---|---|---|---|---|---|
| | Off | On | Off | On | B | Br | U | B | Br | U | Off | On |
| LiGaMD | 2.65 ± 0.34 | 1.59 ± 0.42 | 1.37 ± 0.30 | 2.09 ± 0.19 | 2.23 ± 0.02 | 0.07 ± 0.05 | 0.12 ± 0.00 | 2.42 ± 0.02 | 0.06 ± 0.03 | 0.14 ± 0.01 | 0.94 | 1.28 |
| LiGaMD_Dual | 2.83 ± 0.24 | 1.35 ± 0.25 | 1.82 ± 0.16 | 2.21 ± 0.11 | 2.24 ± 0.00 | 0.07 ± 0.07 | 0.03 ± 0.00 | 2.82 ± 0.03 | 0.06 ± 0.01 | 0.12 ± 0.04 | 0.98 | 1.11 |

(C) CD (q4MD) – 1-Butanol

**Table S4** The ligand binding and unbinding time periods ($\tau_B$ and $\tau_U$) recorded from LiGaMD_Dual simulations of the trypsin-benzamidine binding system.

| System | ID | $\tau_B$ (ns) | $\tau_U$ (ns) |
|---|---|---|---|
| Trypsin - BEN | Sim1 | 14.90, 8.20, 28.90 | 165.90, 77.00, 93.40 |
| | Sim2 | 26.10, 14.00, 27.00, 18.80, 124.10 | 11.50, 270.70, 70.20, 415.70 |
| | Sim3 | 47.39, 31.59, 12.63, 18.29, 49.09, 19.84, 41.00, 15.00, 85.13, 17.19, 60.59 | 145.08, 17.21, 18.71, 28.50, 38.37, 18.52, 32.75, 205.65, 13.37, 32.84 |
| | Sim4 | 32.10, 70.00, 38.70, 30.30 | 210.40, 9.40, 21.70, 496.90 |
| | Sim5 | 7.00, 46.70, 31.30, 39.60 | 279.80, 179.30, 265.80 |

**Table S5** Energy barriers of trypsin-benzamidine dissociation ("off") and binding ("on") calculated from the reweighed ($\Delta F$) and modified (no reweighting, $\Delta F^*$) free energy profiles, curvatures of the reweighed ($w$) and modified ($w^*$) free energy profiles near the guest Bound ("B"), Barrier ("Br") and Unbound ("U") states, and the ratio of apparent diffusion coefficients calculated from the LiGaMD_Dual simulations without reweighting (modified, $D^*$) and with reweighting ($D$).

| Sim | $\Delta F$ (kcal/mol) | | $\Delta F^*$ (kcal/mol) | | $w$ | | | $w^*$ | | | $D^*/D$ | |
|---|---|---|---|---|---|---|---|---|---|---|---|---|
| | Off | On | Off | On | B | Br | U | B | Br | U | Off | On |
| LiGaMD_Dual | 12.17 ± 1.54 | 3.04 ± 2.04 | 1.37 ± 0.56 | 2.40 ± 0.41 | 2.39 ± 0.21 | 0.12 ± 0.16 | 0.06 ± 0.01 | 0.99 ± 0.05 | 0.04 ± 0.05 | 0.06 ± 0.02 | 1.06 | 15.07 |

**Figure S1** Time courses of host-guest distances calculated from (A) LiGaMD and (B) LiGaMD_Dual simulations of CD using the GAFF force field with aspirin, (C) LiGaMD and (D) LiGaMD_Dual simulations of CD using the GAFF force field with 1-butanol, (E) LiGaMD and (F) LiGaMD_Dual simulations of CD using the q4MD force field with aspirin, (G) LiGaMD and (H) LiGaMD_Dual simulations of CD using the q4MD force field with 1-butanol.

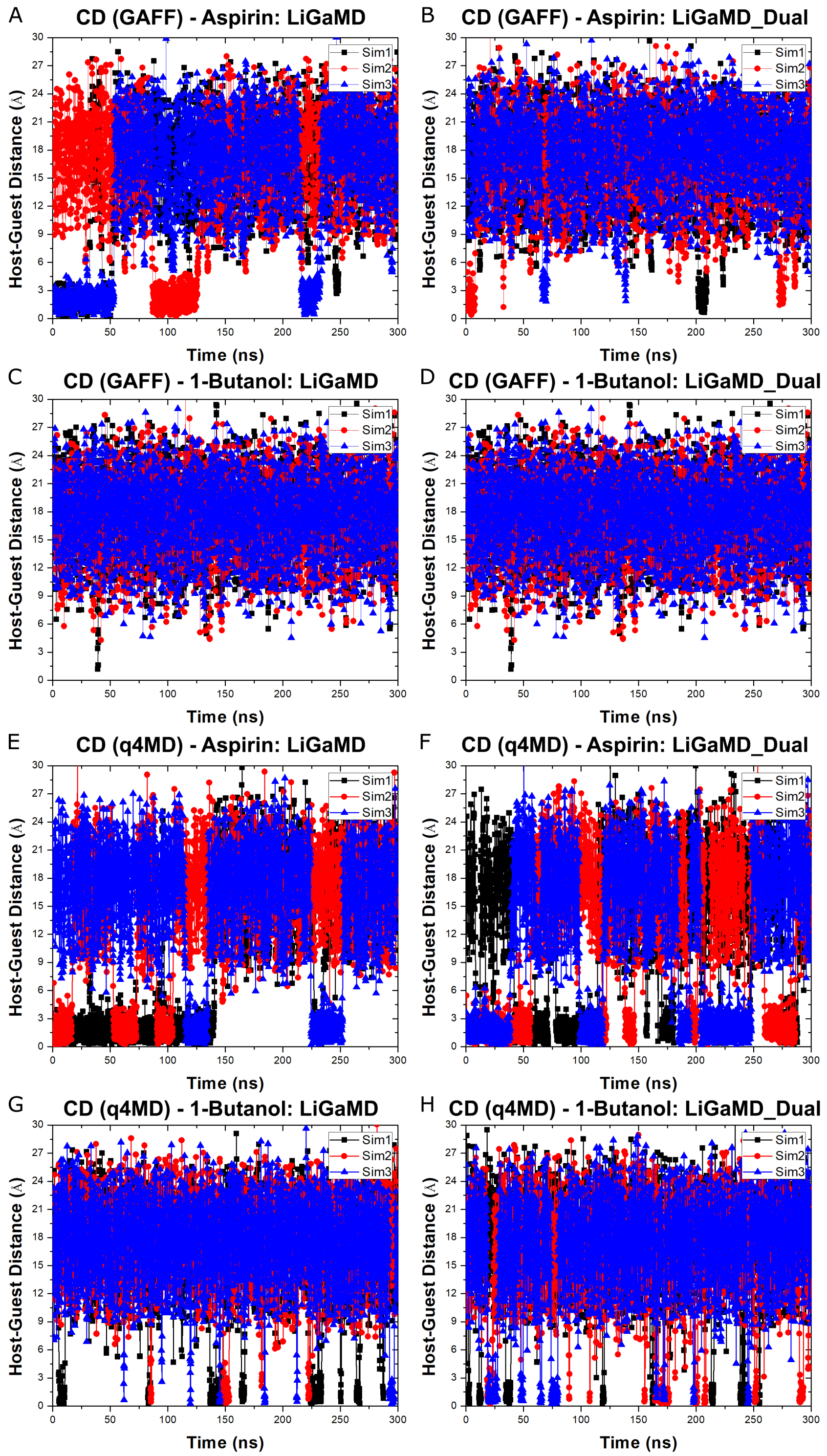

**Figure S2** Reweighted and modified PMF profiles of guest 1-butanol binding to the CD host modeled with the GAFF force field: (A) LiGaMD and (B) LiGaMD_Dual simulations.

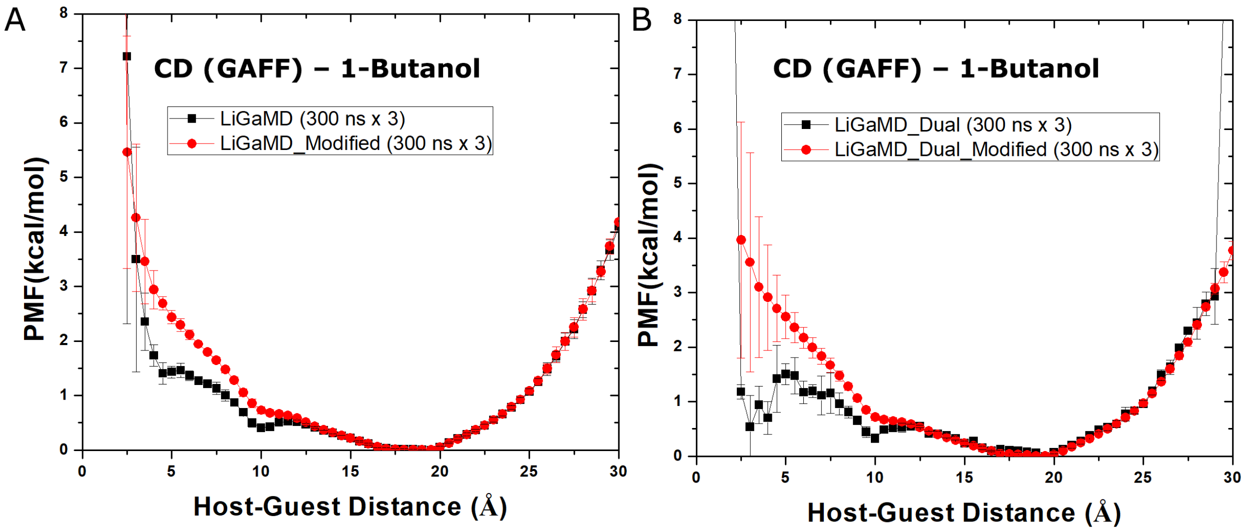

**Figure S3** Time courses of the benzamidine ligand RMSD relative to the X-ray conformation obtained from LiGaMD_Dual equilibration simulations of trypsin, where the input parameter $\sigma_{0P}$ was increased from 1.0 to 6.0 with the threshold energy set to upper bound for applying boost potential to the ligand non-bounded potential energy.

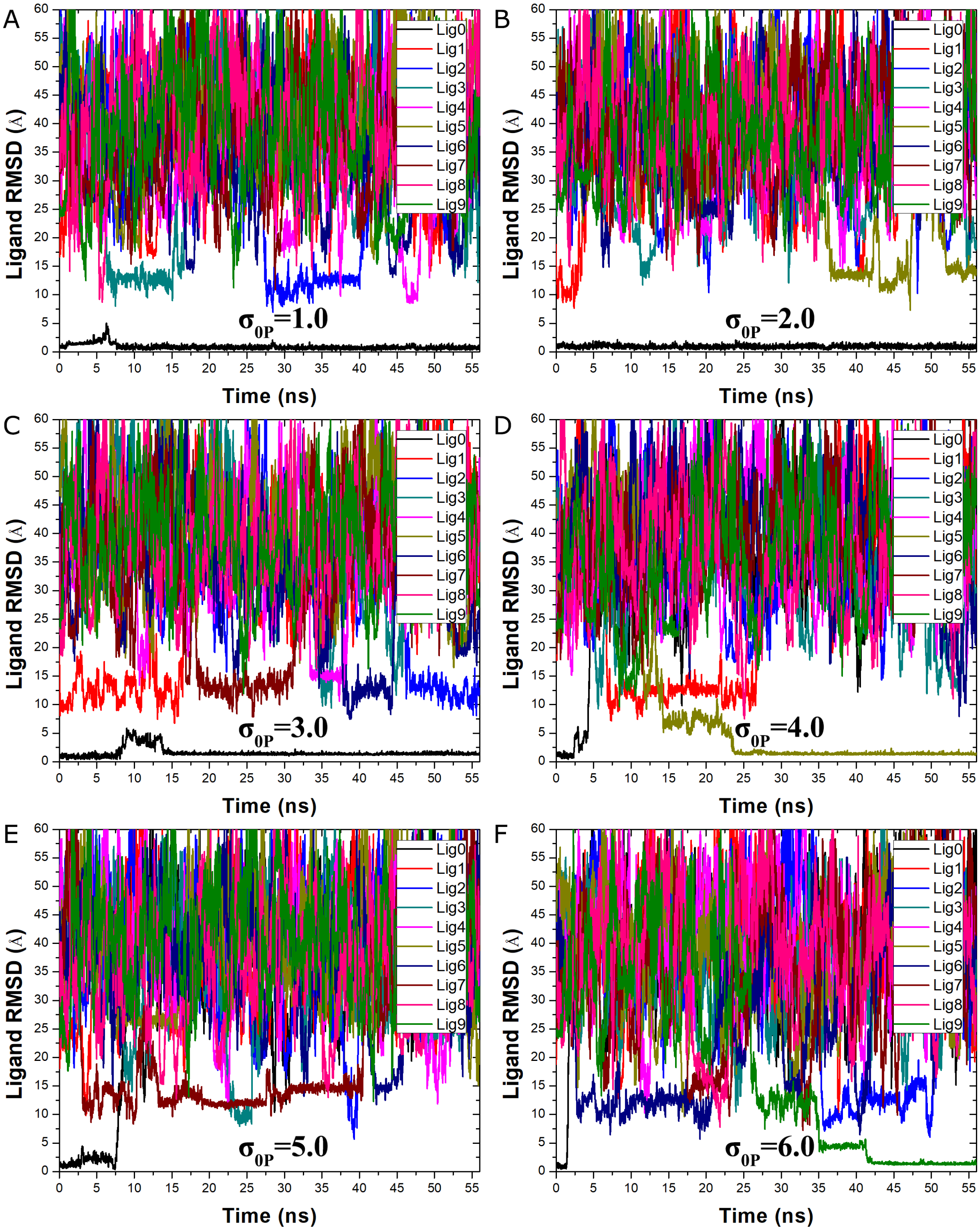

**Figure S4** RMSD of the benzamidine (BEN) ligand relative to the X-ray crystal conformation calculated from LiGaMD_Dual simulations as listed in **Table 4**: (A) Sim1, (B) Sim2, (C) Sim3, (D) Sim4 and (E) Sim5.

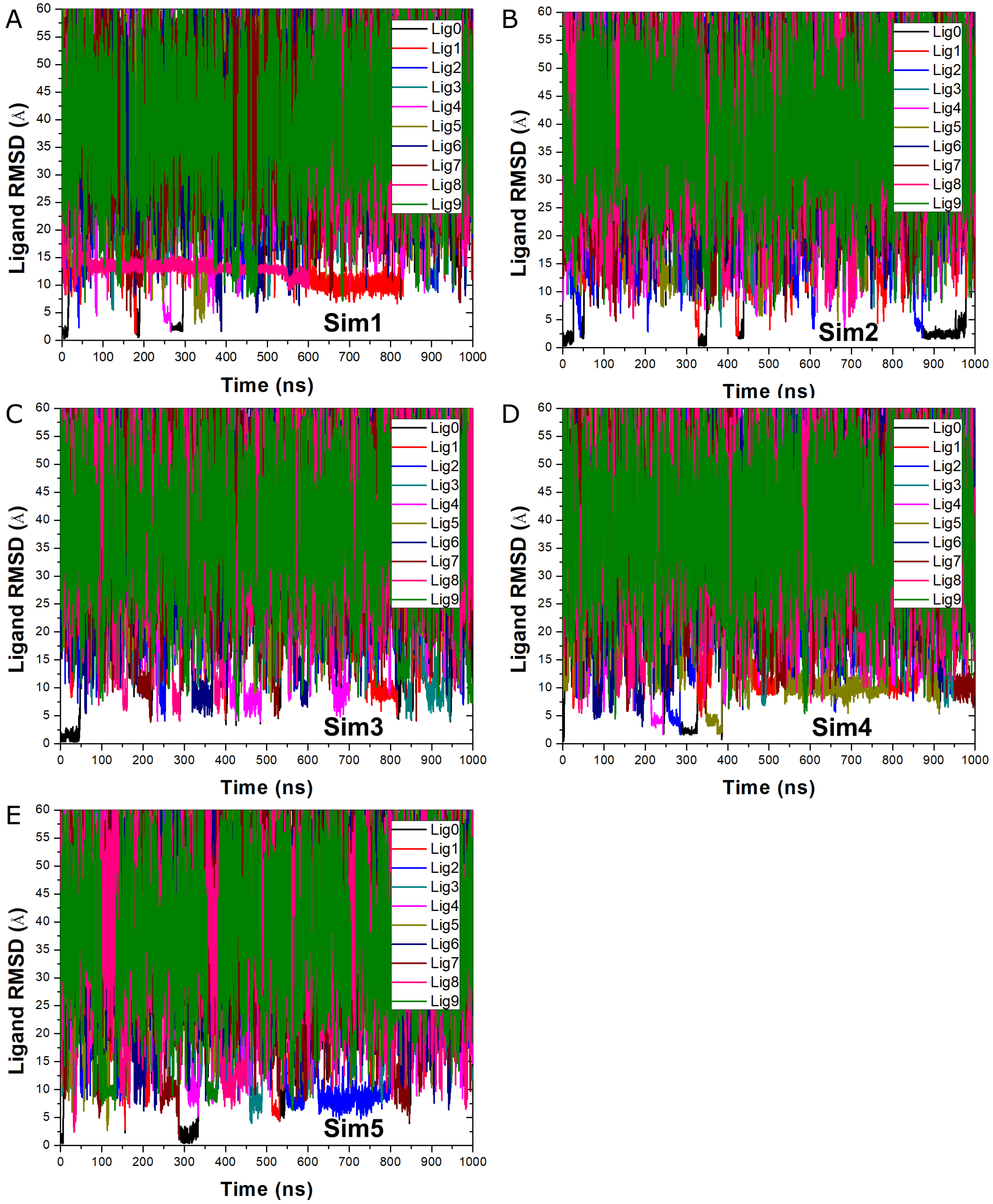

**Figure S5** Distances between the N atom in benzamidine and CG atom of Asp189 in trypsin calculated from five LiGaMD_Dual simulations as listed in **Table 4**: (A) Sim1, (B) Sim2, (C) Sim3, (D) Sim4 and (E) Sim5. Distance plots of Sim2 are provided in **Figure 5A**.

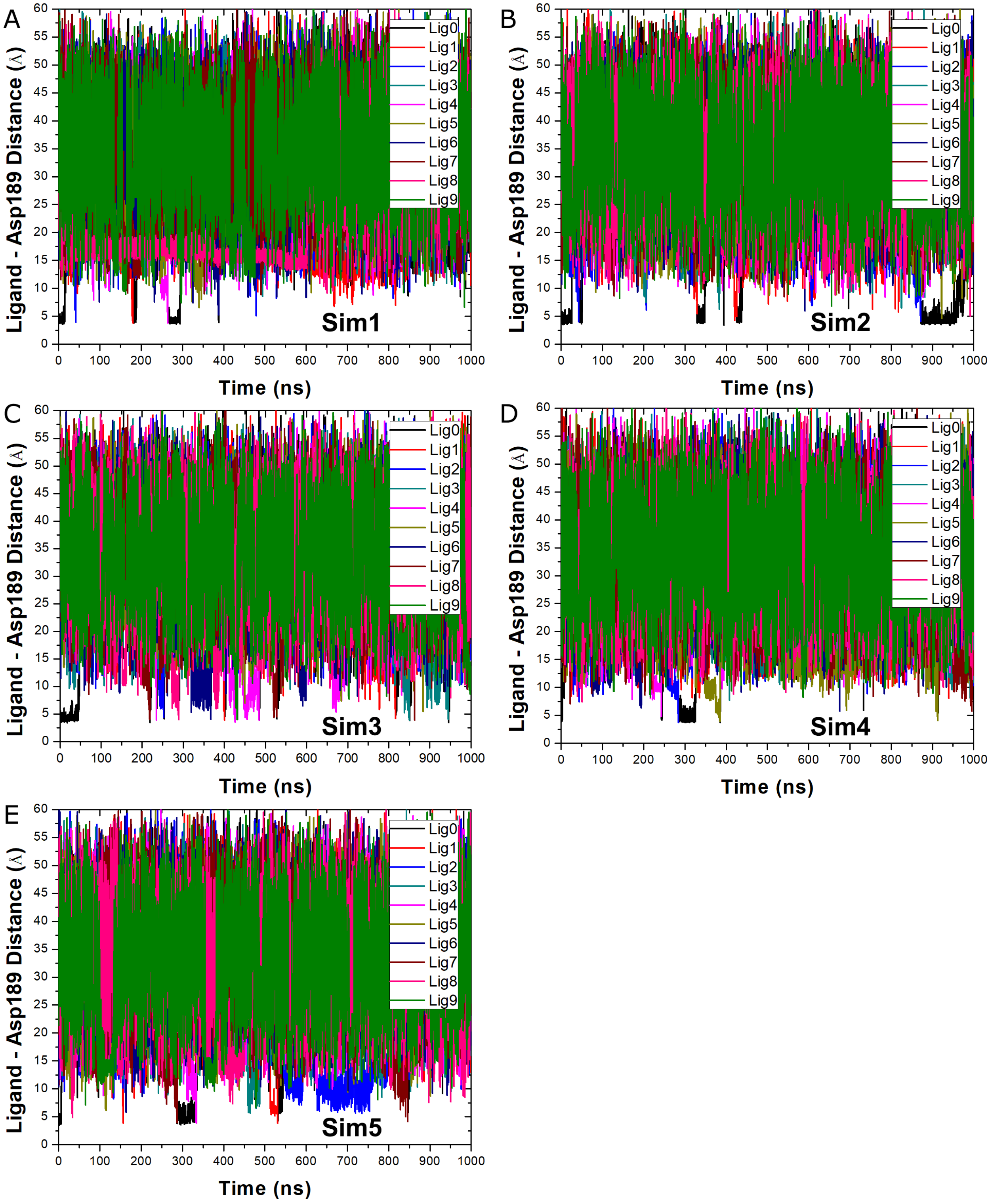

**Figure S6** Reweighted 2D PMF profiles of the BEN:N – Asp189:CG and Trp215:NE – Asp189:CG atom distances calculated from five individual 1000 ns LiGaMD_Dual simulations of the benzamidine inhibitor binding to trypsin: (A) Sim1, (B) Sim2, (C) Sim3, (D) Sim4 and (E) Sim5.

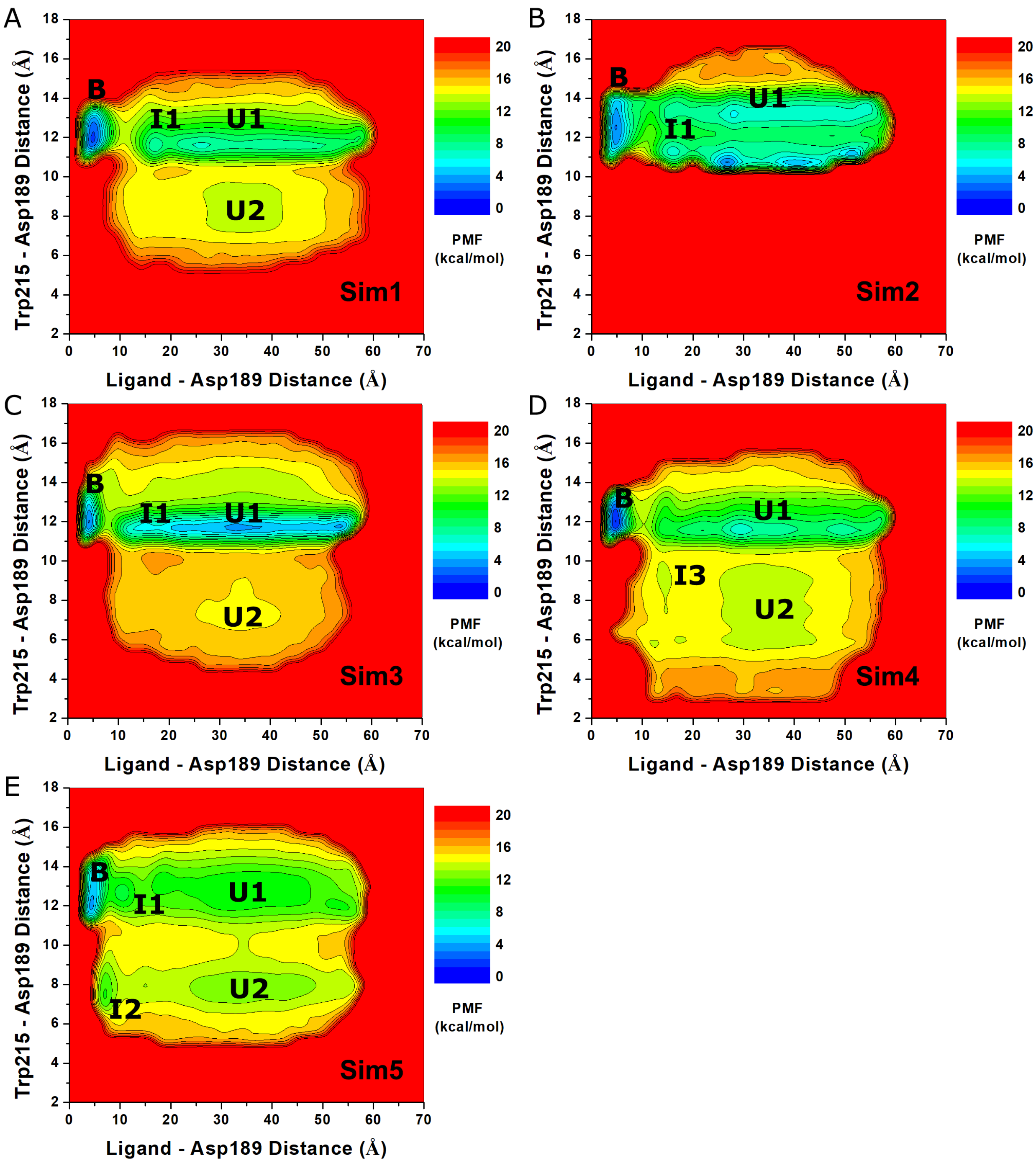

**Figure S7** Modified 2D PMF profiles of the BEN:N – Asp189:CG and Trp215:NE – Asp189:CG atom distances calculated from five individual 1000 ns LiGaMD_Dual simulations of the benzamidine inhibitor binding to trypsin: (A) Sim1, (B) Sim2, (C) Sim3, (D) Sim4 and (E) Sim5.

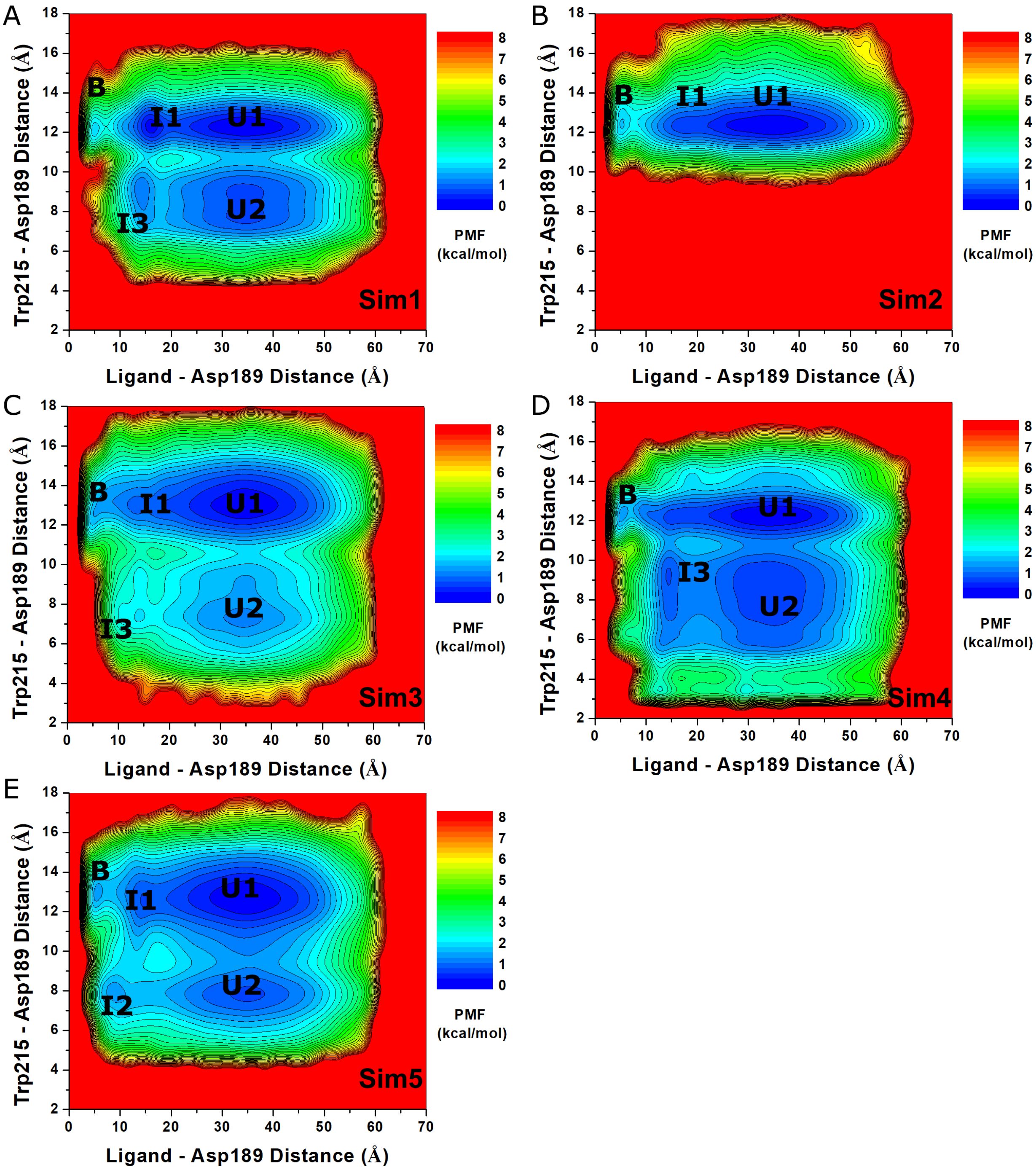
